# The Wnt/APC destruction complex targets SREBP2 in a β-catenin-independent pathway to control cholesterol metabolism

**DOI:** 10.1101/2025.10.12.681896

**Authors:** Ahmed Rattani, Cora Anderson, William V. Trim, Esteban Garita, Shinya Imada, Saleh Khawaled, Heaji Shin, Jiuchun Zhang, Ömer H. Yilmaz, Marc Kirschner

**Affiliations:** Department of Systems Biology, Harvard Medical School, Boston, MA 02115, USA; Division of Hematology/Oncology, Department of Medicine and Cancer Research Institute, Beth Israel Deaconess Medical Center, Harvard Medical School, Boston, MA 02215, USA; Department of Biology, The David H. Koch Institute for Integrative Cancer Research at MIT, MIT, Cambridge, MA 02139, USA; Department of Cell Biology, Harvard Medical School, Boston, MA, 02115; Department of Pathology, Beth Israel Deaconess Medical Center, Harvard Medical School, Boston, MA 02215, USA

## Abstract

APC, the core scaffold of the Wnt destruction complex, targets the transcriptional co-activator β-catenin for proteolysis. There is no convincing evidence that APC directs degradation of other substrates. Using a reconstituted cytosolic extract-based system and complementary in vivo and cellular assays, we show that SREBP2, the master regulator of cholesterol biosynthesis, is a direct APC–AXIN1 substrate. APC-dependent SREBP2 degradation is conserved in *Xenopus* embryos, mouse colon, and human colorectal cancer cells and restricts SREBP2 target-gene expression, cholesterol synthesis, and tissue cholesterol levels. Mechanistically, APC and AXIN1 promote SREBP2 degradation via a conserved phosphodegron, which marks SREBP2 for ubiquitination by the E3 enzyme, FBXW7. Like β-catenin, SREBP2 is stabilized by extracellular Wnt ligands; unlike β-catenin, its regulation is independent of GSK3β and CK1α and requires the entire APC mutational cluster region (MCR), whereas β-catenin turnover can operate with only a partial MCR. These findings define a β-catenin-independent branch of Wnt signaling that couples APC to sterol metabolism, providing a mechanistic rationale to target the mevalonate/SREBP2 axis in APC-mutant colorectal cancer.

## INTRODUCTION

The canonical Wnt/β-catenin pathway is a highly conserved signaling cascade essential in multiple steps in metazoan development and for adult tissue homeostasis (Nusse & Clevers, 2017). The pathway is governed by a multi-protein “destruction complex”-comprised of the scaffold proteins Adenomatous polyposis coli (APC) and AXIN1 alongside the kinases glycogen synthase kinase 3 (GSK3) and casein kinase 1 (CK1) (Rubinfeld et al., 1993; Zeng et al., 1997, Ranes, et al., 2021). These kinases sequentially phosphorylate the transcriptional co-activator β-catenin, targeting it for proteasomal degradation (Amit et al., 2002; Liu et al., 2002; Yost et al., 1996, Ranes et al., 2021, Hernández et al., 2012). Signaling in the Wnt pathway occurs when the WNT ligand binds to the transmembrane Frizzled (FZD) receptors and co-receptors, LRP5/6, leading to the inactivation of the APC and AXIN complex, thereby blocking the constitutive degradation of β-catenin. The resulting rise in β-catenin drives a transcriptional program that controls cell fate, cell proliferation, and tissue patterning (Molenaar et al., 1996) in many biological contexts found in virtually all metazoans (Bhanot et al., 1996; Pinson et al., 2000; Tamai et al., 2000). Pathway dysregulation can have profound consequences; its hyperactivation by loss-of-function mutations in APC, acting as a tumor suppressor, is a well-established driver of colorectal cancers (Kinzler & Vogelstein, 1996; Zhan et al., 2017).

Despite its well understood framework, a central question persists: are the diverse effects of the destruction complex in so many biological contexts always mediated solely through β-catenin, or does the WNT destruction complex also regulate other targets important in cell differentiation, stem-cell behavior and tumor initiation? Paradoxically, in colorectal cancer where inactivating mutations in APC are very common (∼80% of cases), activating mutations in β-catenin (CTNNB1) are very rare (Markowitz & Bertagnolli, 2009), perhaps indicating a much broader (and unappreciated) regulatory role for APC than for β-catenin. Although there is no direct evidence to suggest other targets, it has seemed possible that APC possesses other functions beyond its ability to mediate the degradation of β-catenin (Näthke, 2004).

Uncovering other functions of APC has been experimentally difficult. Conventional genetic approaches, such as deleting APC or AXIN1, are confounded because the loss of these core destruction-complex components inevitably stabilizes β-catenin, which could lead to pleiotropic downstream transcriptional effects (Funayama et al., 1995; Yost et al., 1996). To circumvent these limitations, we looked upstream of potential transcriptional processes by directly probing the immediate reactions in the destruction complex, developing a nucleus-free cytosolic extract based in vitro system that reconstitutes APC-mediated β-catenin degradation independent of any possible transcriptional input.

Using this approach, we identified the master transcriptional regulator of cholesterol metabolism, sterol regulatory element-binding protein 2 (SREBP2) as a direct target of the APC/AXIN1 destruction complex, previously thought to exclusively regulate β-catenin stability. We demonstrated further that APC licenses the E3 (ubiquitin protein ligase) FBXW7 to mediate the degradation of SREBP2, a pathway completely independent of the GSK3β/CK1α-dependent degradation of β-catenin. Both β-catenin and SREBP2 pathways are sensitive to APC truncation; however, when we reconstituted with APC variants commonly observed in human colorectal cancer, many truncated APCs retained substantial β-catenin-degrading activity yet were completely impaired in degrading SREBP2. Such findings define a β-catenin-independent branch of Wnt signaling that directly links the APC tumor suppressor to metabolic control, offering a possible mechanistic explanation for why many colorectal cancers have selected specifically for truncations that abolish the SREBP2 degradation function of APC.

## RESULTS

### A lysate-based assay that recapitulates key features of the WNT destruction complex

After almost 40 years of investigation, β-catenin remains the sole broadly accepted, in vivo validated substrate of APC and the WNT destruction complex; no other protein has been convincingly shown to depend on the APC–AXIN1 scaffold for its turnover. Yet we can still speculate that the diverse biological targets and broad biological functions of this pathway might involve undiscovered direct targets of Wnt regulated protein degradation. With the goal of identifying such novel targets of the WNT destruction complex, should they exist, we developed an in vitro, lysate-based degradation assay. In this system, radiolabeled proteins (putative substrates) were added either to extracts from AXIN1- or APC-deficient cells or to the same extracts supplemented with purified AXIN1 or APC proteins (Fig. 1a, Extended Data Figure 1a–c). We optimized such an assay using β-catenin, the only known canonical substrate of the Wnt mediated destruction complex. As expected, β-catenin showed minimal degradation in the absence of APC or AXIN1 but was efficiently degraded when purified APC or AXIN1 was reintroduced into the corresponding knockout extracts (Fig. 1b, and Extended Data Fig. 1d&e). Thus, the in vitro system faithfully recapitulated the requirement for the core scaffold proteins APC and AXIN1 for β-catenin degradation, consistent with in vivo studies.

**Figure 1.**
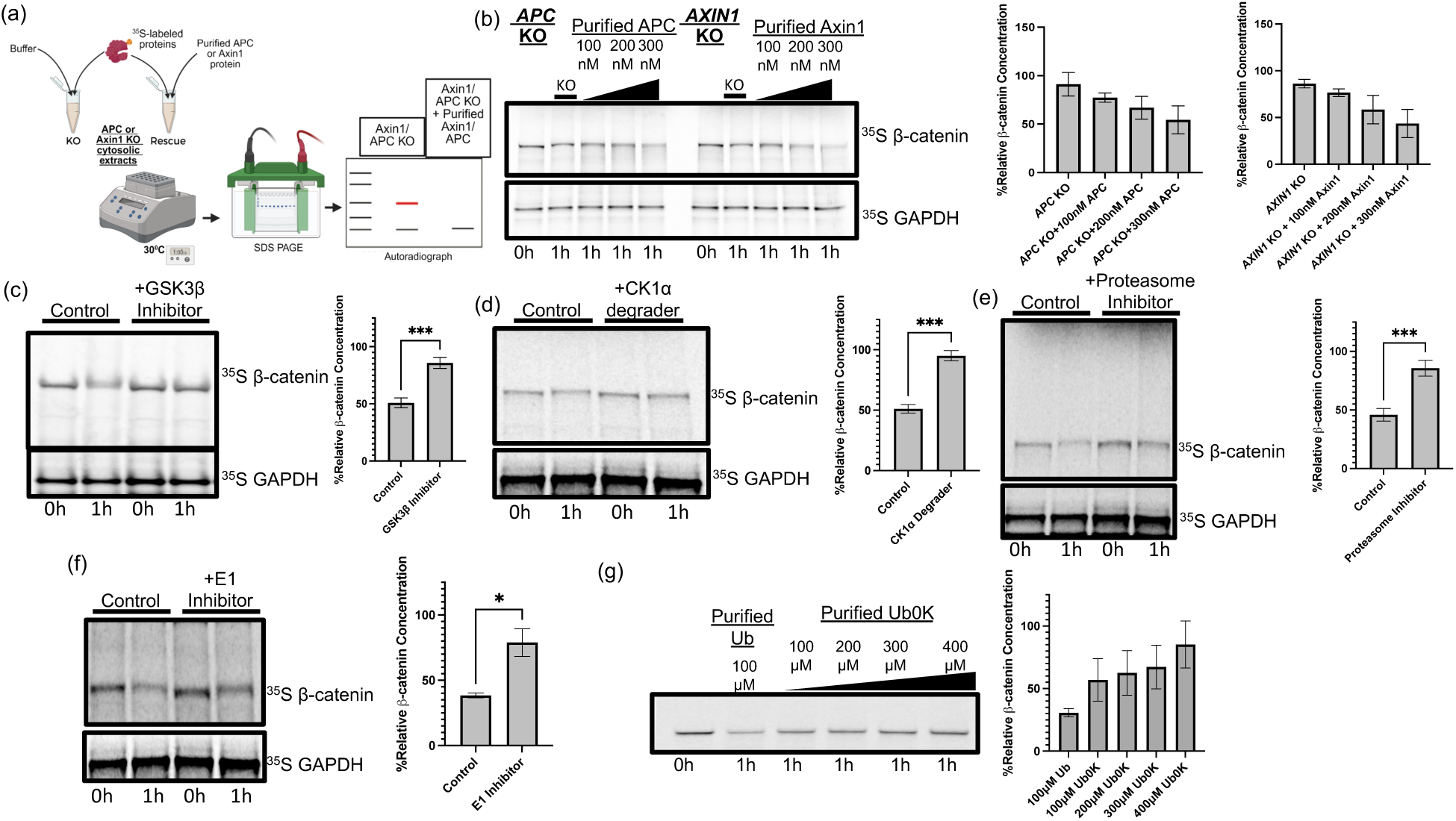
A cell-free system recapitulates the degradation complex mediated degradation of β-catenin. **a**, Schematic of the in-vitro degradation assay. ^35^S-labelled substrate proteins were incubated in nucleus-free cytosolic extracts prepared from wild-type (WT), *AXIN1* knockout (KO) or *APC* KO cells. KO extracts were supplemented with buffer or the corresponding recombinant rescue protein (APC or AXIN1) to restore destruction-complex activity. Reactions were analyzed by SDS–PAGE/autoradiography; substrate stability is quantified relative to t0. **b,** Dose-dependent rescue of β-catenin degradation by APC or AXIN1. Purified APC (left) or AXIN1 (right) was added to the respective knockout extracts. Representative autoradiographs and densitometry are shown. One-way ANOVA revealed significant effects of APC (F(3,8)=5.64, p=0.023) and AXIN1 (F(3,8)=8.78, p=0.0065), and linear regression confirmed dose-dependent reductions in β-catenin (APC: p=0.001, R²=0.68; AXIN1: p=0.00023, R²=0.76). **c, d,** Canonical kinase dependence. Extracts from cells pre-treated with the GSK3β inhibitor LY2090314 (2 μM, 4h) (c) or the CK1α degrader SJ3149 (2 μM, 4 h) (d) stabilize β-catenin and produce an electrophoretic mobility shift consistent with loss of phosphorylation. **e,** β-catenin turnover in cellular extract is proteasome-dependent. Extracts prepared from cells pre-treated with the carfilzomib (10 µM, 4h) resulted in β-catenin stabilization. **f,** Degradation requires the ubiquitin-activating enzyme E1. Pre-treatment with the E1 inhibitor TAK-243 (5 μM, 1h) blocks β-catenin turnover. **g,** Polyubiquitin requirement. Supplementation with wild-type ubiquitin supports degradation, whereas lysine-free ubiquitin (UBK0) prevented turnover and revealed a characteristic ubiquitin ladder on longer exposure. One-way ANOVA indicated a significant effect of Ub/Ub0K concentration (F(4,10)=4.60, p=0.023), and linear regression confirmed a dose-dependent increase in β-catenin (p=0.00081, R²=0.59). Data in **b–g** are mean ± s.d. from n ≥3 independent experiments; representative autoradiographs are shown. (b&e) One-way analysis of variance (ANOVA) with linear regression; (**c-e**) Unpaired two-tailed *t*-test.

We further validated our assays by confirming that β-catenin degradation requires phosphorylation by GSK3β and CK1α, using specific inhibitors for GSK3β, degrader treatment for CK1α, and mutant β-catenin lacking phosphorylation sites. These perturbations not only reduced β-catenin degradation but also increased protein mobility on gels, consistent with the expected loss of phosphorylation (Fig. 1c&d, Extended Data Figure 1e). In addition, proteasome inhibition, blockade of the E1 ubiquitin-activating enzyme, and expression of lysine-free ubiquitin each prevented β-catenin degradation (Fig. 1e-g). Prolonged exposure of radiolabeled gels from the lysine-free ubiquitin experiments revealed a characteristic polyubiquitination ladder, further supporting ubiquitin-dependent turnover of β-catenin in the cytosolic extract, as in the cells (Extended Data Fig. 1f).

Together, these results demonstrate that the lysate-based system preserves the essential biochemical features of the endogenous WNT destruction complex, providing a tractable platform to identify and test novel substrates.

### SREBP2 is a conserved substrate of the APC-Axin1 destruction complex

To identify new substrates of the WNT destruction complex, we applied our in vitro degradation assay to a library of human transcription factors and identified SREBP2, a key regulator of the cholesterol homeostasis pathway (Wang et al., 1993; Brown & Goldstein, 1997; Horton et al., 2002), as a previously unrecognized target of APC (Extended Data Fig. 2a). When purified APC protein was added, degradation of ^35^S-labeled SREBP2 commenced in a dose-dependent manner, whereas AXIN1 alone exerted only a modest effect (about 50% degradation of SREBP2 noted with 300nM APC compared to less than 30% with 300nM Axin1) (Fig. 2a). Notably, APC failed to promote SREBP2 degradation in extracts lacking AXIN1; however, reintroduction of low amounts of AXIN1 to AXIN1-deficient extracts (AXIN1 was added at 100nM, slightly below the previously reported physiological range for the HEK293T cells (Tan, C. W. et. al. 2012)) restored APC-dependent degradation of SREBP2 in a dose-dependent manner (Fig. 2b&c). These findings establish that both APC and AXIN1 are both required for SREBP2 turnover in this cytosolic extract-based system. As expected, SREBP2 turnover in cell extracts required the ubiquitin-proteasome system, as the proteasome inhibitor stabilized SREBP2, and titration of lysine-free ubiquitin (UBK0) progressively blocked degradation and revealed a characteristic ubiquitin ladder (Extended Data Fig. 2b&c).

**Figure 2.**
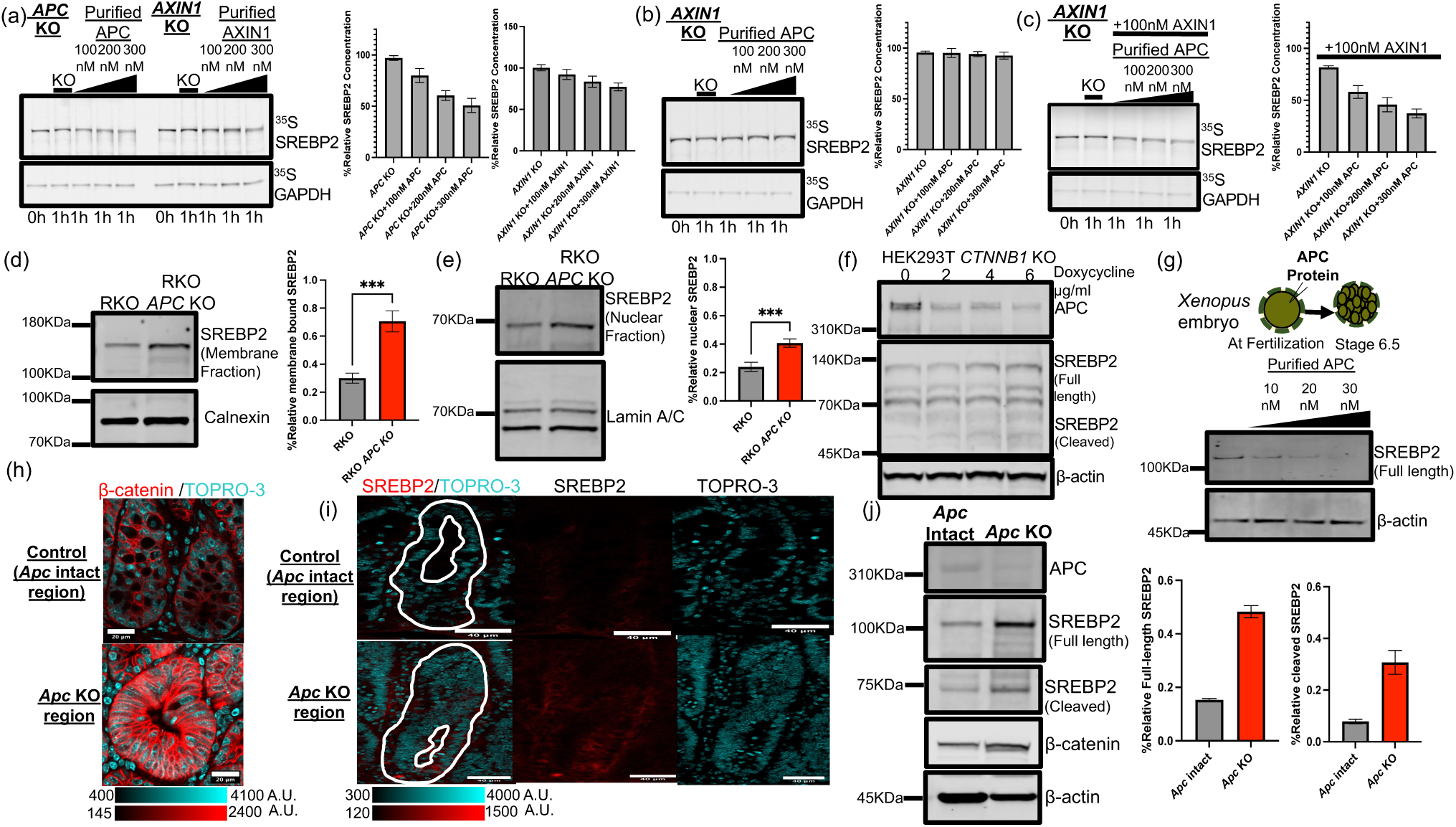
SREBP2 is a conserved, β-catenin-independent substrate of the APC–AXIN1 destruction-complex. **a**, APC drives dose-dependent SREBP2 degradation in vitro. ³⁵S-labeled SREBP2 was incubated in *APC*/*AXIN1* KO lysate supplemented with increasing concentrations of purified reconstituted protein. ANOVA showed significant APC and AXIN1 dose effects (p<0.0001). Regression confirmed dose-dependent SREBP2 degradation (APC: −0.158, R²=0.92; AXIN1: −0.078, R²=0.76; p<0.0001). **b,** AXIN1 is required for APC-mediated turnover. Adding recombinant APC to *AXIN1*-KO lysate does not promote SREBP2 degradation. One-way ANOVA showed no significant dose effect (F(3,8)=0.53, p=0.676), and linear regression confirmed the absence of a dose-dependent trend (slope = −0.010, p=0.207, R²=0.15). **c,** AXIN1 reconstitution restores APC activity. Low-dose AXIN1 added to *AXIN1*-KO lysate rescues APC-dependent, dose-responsive SREBP2 degradation (F(3,8)=42.6, p<0.0001; slope −0.146, p<0.0001, R²=0.89). **d,** *APC* loss increases membrane-associated SREBP2. Subcellular fractionation of cholesterol-deprived RKO and RKO APC-KO cells shows increased SREBP2 in the membrane fraction (0.294 ± 0.034 vs. 0.710 ± 0.070; t=−9.74, p<0.0001). **e,** *APC* loss increases nuclear SREBP2. Subcellular fractionation of cholesterol-deprived RKO and RKO *APC* KO cells shows increased SREBP2 in the nuclear fraction (0.241 ± 0.029 vs. 0.406 ± 0.028; t=−7.73, p<0.0001). **f,** APC regulates SREBP2 independently of β-catenin. Immunoblot of β-catenin-null cells bearing a doxycycline-inducible APC shRNA. Increasing doxycycline causes a dose-dependent reduction in APC with a reciprocal increase in SREBP2. **g,** APC regulates SREBP2 post-translationally. Injection of purified human APC protein into single-cell *Xenopus* embryos (pre–mid-blastula, transcriptionally silent) reduces endogenous SREBP2 dose-dependently. Texas Red dye was used to confirm micro-injection. **h,** Focal *Apc* deletion drives nuclear β-catenin. In *Apc^fl/f^;Villin^CreERT2^* colon, β-catenin immunostaining shows increased nuclear β-catenin and was used to confirm *Apc* knockout in mutant crypts relative to matched adjacent control. Scale bar, 20 μm. **i,** Focal epithelial *Apc* deletion increases SREBP2 relative to the matched adjacent control from the same animal. Immunofluorescence of *Apc^fl/fl^;Villin^CreERT2^* colon shows SREBP2 (red) with TO-PRO-3 nuclear counterstain (cyan). Scale bar, 40 μm. **j,** Focal epithelial Apc deletion increases full-length and cleaved SREBP2. Immunoblot of *Apc^fl/fl^;Villin^CreERT2^* colon shows increase in full-length (0.153 ± 0.005 vs. 0.482 ± 0.023; t=−24.45, p<0.001), and cleaved SREBP2 (0.079 ± 0.009 vs. 0.307 ± 0.046; t=−8.42, p<0.02) in focal *Apc*-deleted tissue relative to the matched adjacent control from the same animal. *For **a–e**, j, representative autoradiographs/immunoblots are shown with quantification on right. Data are mean ± s.d. from ≥3 independent experiments*.

We show that the in vitro assays were consistent with studies of SREBP2 regulation in cells. SREBP2 levels were elevated in APC-deficient cells compared with that of controls (Extended Data Fig. 2d&e). Furthermore, subcellular fractionation showed increased SREBP2 abundance in both membrane and nuclear compartments (Fig. 2d, e). Short hairpin RNA (shRNA)-mediated depletion of *Apc* further increased SREBP2 levels in a dose-dependent manner in both β-catenin-intact colorectal cancer cells (HCT116) and β-catenin-knockout HEK293T cells, further demonstrating that APC regulates SREBP2 independently of β-catenin (Fig. 2f; Extended Data Figure 2f).

Consistent with these findings, cycloheximide-chase assays showed a slower rate of SREBP2 degradation in APC-knockout cells than in controls (Extended Data Figure 2g). Moreover, GFP pull-down from endogenously tagged APC-GFP lines confirmed that APC and SREBP2 interact (Extended Data Fig. 2h).

To definitively exclude the possibility that APC regulates SREBP2 through transcriptional feedback, we microinjected purified APC protein into single-cell *Xenopus* embryos, which lack transcription until the mid-blastula transition at the 4000-cell stage (Newport & Kirschner, 1982). Under these conditions, we observed a dose-dependent reduction in SREBP2 levels, providing strong evidence that APC regulates SREBP2 through an evolutionarily conserved post-translational mechanism (Fig. 2g).

To validate these findings in vivo, we used a *Apc^f/f^;Villin^Cre-ERT2^* mouse model in which colon adenomas could be induced by colonoscopy guided 4-hydroxy-tamoxifen injections into the intestinal epithelium to induce focal *Apc* deletion, leaving the adjacent colon with intact *Apc* as an internal control (Fig. 2h) (Roper et al., 2017; Roper et al., 2018), and therefore minimizing inter-animal and dietary variabilities. In the transformed tumor area with the deletion of *Apc*, we observed increased levels of both full-length and cleaved SREBP2 relative to the matched control regions (Fig. 2i, j). Together, these results demonstrate that APC directly regulates SREBP2 stability in colonic epithelial cells.

### APC deletion enhances SREBP2 target gene expression and cholesterol synthesis

To evaluate the functional consequences of SREBP2 stabilization, we measured the expression of canonical SREBP2 target genes in *Apc*-deficient cell lines and tissues. Consistent with increased SREBP2 activity, *Apc* deletion led to elevated mRNA expression of cholesterol biosynthetic targets when compared with matched *Apc*-intact regions from the same mice (Fig. 3a). At the protein level, canonical SREBP2 targets regulated by APC, include: 3-hydroxy-3-methylglutaryl-CoA reductase (HMGCR; the rate-limiting enzyme in cholesterol biosynthesis), 3-hydroxy-3-methylglutaryl-CoA synthase 1 (HMGCS1; catalyzes formation of HMG-CoA from acetoacetyl-CoA), low-density lipoprotein receptor (LDLR; mediates cholesterol uptake), and farnesyl diphosphate synthase (FDPS; generates farnesyl diphosphate for downstream sterol synthesis), were each increased (Horton et al., 2002; Horton et al., 2003). In contrast, farnesyl-diphosphate farnesyltransferase 1 (FDFT1; squalene synthase, the first committed enzyme in the sterol branch of the mevalonate pathway) and mevalonate kinase (MVK, which catalyzes the first step after mevalonate formation) that have not been demonstrated to be direct SREBP2 targets, were unchanged relative to controls in mice (Fig. 3b). In RKO cells, APC KO increased proteins levels all downstream targets, except FDPS (Extended Data. Fig. 3).

**Figure 3.**
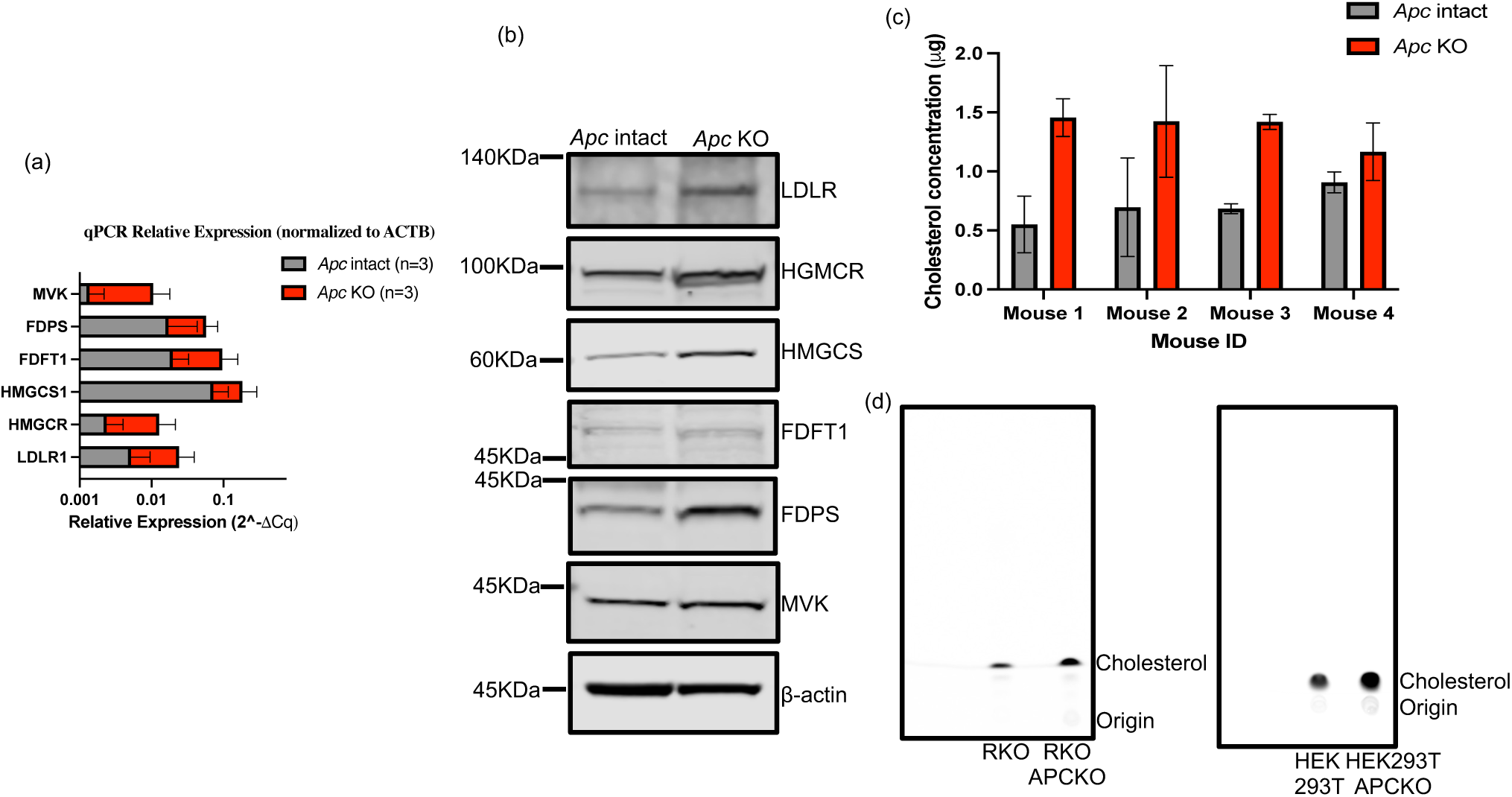
APC regulates SREBP2 downstream targets and cholesterol synthesis. **a**, Upregulation of SREBP2 target gene transcripts in *Apc*-deficient colon. RT-qPCR analysis shows increased transcripts for key SREBP2 target genes (including enzymes for cholesterol synthesis) in epithelial cells from *Apc*-deficient colonic tissue compared to adjacent wild-type (WT) tissue from the same mice. Data are *mean ± s.d.* (n=3 mice). **b,** Increased SREBP2 target protein levels in *Apc*-deficient colon. A representative Western blot confirms that the transcriptional changes lead to increased protein levels of key SREBP2 target genes in *Apc*-deficient tissue relative to matched WT controls. β-actin serves as a loading control. **c,** Elevated cholesterol content in *Apc*-deficient colonic epithelium. Direct quantification of total cholesterol in isolated epithelial cells from four individual mice reveals significant accumulation in *Apc*-deficient regions compared to matched WT controls (paired analysis: 0.709 ± 0.145 vs. 1.366 ± 0.141; t=7.30, p=0.005). Data are *mean ± s.d.* of technical replicates. **d,** Increased *de novo* cholesterol synthesis in *APC* KO cells. Control (RKO and HEK293T) and *APC* knockout (RKO *APC* KO and HET293T *APC* KO) cells were sterol-depleted for 16 h, and then pulsed with [1-^14^C] acetate in complete medium. Thin-layer chromatography (TLC) and autoradiography show increased incorporation of the radiolabel into newly synthesized cholesterol in *APC* KO cells.

To assess whether the observed transcriptional changes were accompanied by functional alterations in cholesterol metabolism, we measured total cholesterol levels in transformed tumors using the conditional *Apc^f/f^;Villin^Cre-ERT2^* colon model as described above. The cholesterol content, quantitated by a fluorometric assay, was approximately two-fold higher in *Apc*-deleted intestinal epithelial cells than in matched control regions with intact *Apc* from the same animals (Fig. 3c). We also examined *de novo* cholesterol synthesis after sterol depletion by pulsing with [1-^14^C] acetate, followed by saponification and resolution by thin-layer chromatography; APC-deficient cells incorporated about two-fold more label into cholesterol than APC-intact controls (Fig. 3d&e). Together, these findings clearly indicate that APC loss not only stabilizes SREBP2 but also upregulates its downstream targets and enhances cholesterol biosynthesis in both the colonic epithelium and cell lines.

### SREBP2 is regulated by upstream Wnt signaling independently of β-catenin but requires APC

APC-mediated β-catenin turnover is well known to be controlled by extracellular Wnt ligands (Nusse & Clevers, 2017). Hence, we asked whether SREBP2 responds to the same inputs. HEK293T wild-type and β-catenin-knockout (KO) cells were stimulated with Fzd7/8 subtype NGS Wnt (Wnt Surrogate-Fc) (Janda et al., 2017, Miao et al. 2020) together with R-spondin-1 (to increase the potency of the Wnt signal) (de Lau et al., 2011). We used LRP6 phosphorylation to verify pathway activation. Under such conditions, SREBP2 protein increased over time in both wild-type and β-catenin-KO cells (about 3–5-fold more at 4 h versus t_0_), but not in APC-KO cells, indicating that the response is β-catenin-independent and yet APC-dependent (Fig. 4a, b; Extended Data Fig. 4a). Consistent with their roles in Wnt signaling upstream of the destruction complex, Wnt-induced SREBP2 accumulation was abolished in dishevelled (*DVL*) KO cells and in Frizzled (*FZD*) KO cells (Extended Data Fig. 4b&c).

**Figure 4.**
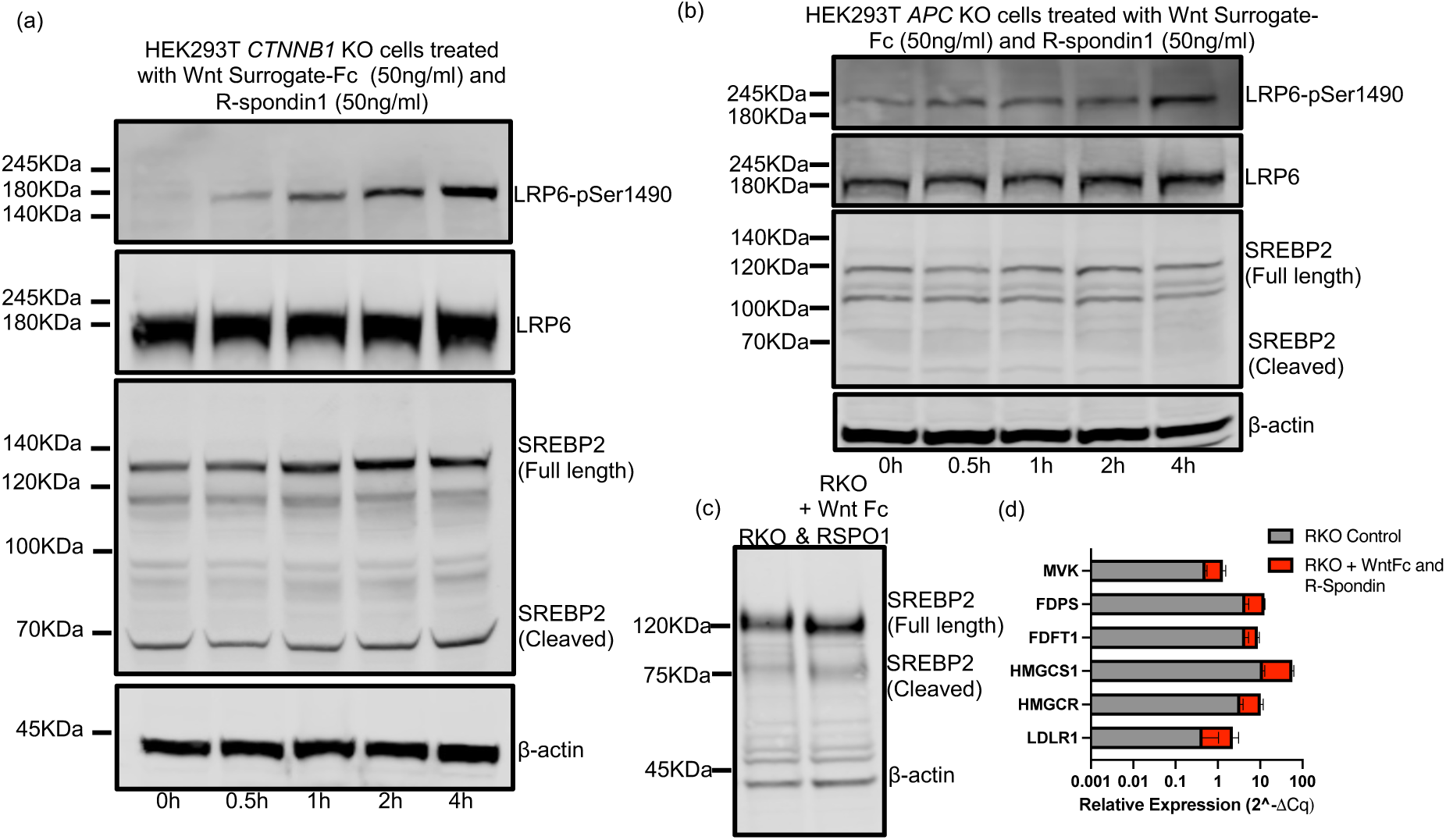
Wnt signaling stabilizes SREBP2 and activates its downstream pathway in an APC-dependent, β-catenin-independent manner. **a**, Western blot analysis of *CTNNB1* (β-catenin) knockout (KO) HEK293T cells. Cells were cholesterol-deprived for 16 h, then shifted to complete medium with Wnt Surrogate-Fc and R-spondin-1 (50 ng/ml each) over a 4-hour time course. Wnt pathway activation, confirmed by LRP6 phosphorylation at Ser1490, leads to the rapid stabilization of SREBP2. **b,** Western blot of *APC* KO HEK293T cells treated as in (a). Wnt stimulation does not alter SREBP2 levels in the absence of APC, indicating that APC is required to mediate the effects of Wnt signaling on SREBP2 stability. **c,** Cells treated as in (a). Western blot showing that treatment of RKO colorectal cancer cells with Wnt Surrogate-Fc and R-spondin-1 for 4 hours also stabilizes SREBP2. **d,** The Wnt-induced stabilization of SREBP2 is functionally active. RT-qPCR analysis of RKO cells treated as in (c) shows a significant upregulation in the transcription of downstream SREBP2 target genes. Data for n=2 samples are shown as *mean ± s.d.*.

We next asked whether Wnt stimulation augments the downstream transcriptional output of the SREBP2 pathway. To stimulate SREBP2 protein cleavage, cells were cultured in sterol-depleted medium and then placed in medium with normal serum in the presence or absence of the Wnt Surrogate-Fc plus R-spondin-1. Wnt treatment increased both precursor (full-length) and nuclear (cleaved) SREBP2 and produced a modest but consistent elevation in canonical SREBP2 target genes relative to controls (Fig. 4c, d). Together, these results show that, similar to β-catenin, SREBP2 is under the control of upstream WNT–FZD–DVL signaling and that its regulation by this pathway operates independently of β-catenin but requires APC, further demonstrating a connection between sterol regulation and APC dependent WNT destruction complex.

### APC-mediated SREBP2 degradation requires phosphorylation of a conserved FBXW7 phosphodegron

The E3 ubiquitin ligase FBXW7 is known to target SREBP2 for proteasomal degradation, but only after phosphorylation within a conserved phosphodegron (CPD) (Sundqvist et al., 2005). Because GSK3β primes several FBXW7 substrates (Welcker & Clurman, 2008), we asked whether phosphorylation is likewise required for the APC-mediated degradation of SREBP2.

In *Xenopus* egg extracts, which we and others have previously shown to maintain both phosphorylation status and protein degradation activity for a prolonged periods of time in the studies of cell cycle substrates (Miake-Lye & Kirschner, 1985; Lohka & Maller, 1985), the nuclear (S2) form of wild-type SREBP2 became strongly phosphorylated.

However, the CPD mutant S432A/S436A showed little to no phosphorylation in *Xenopus* egg extracts, suggesting this motif as the principal site of modification (Fig. 5a). Purified APC (capable of complementing APC deletion confirmed by β-catenin and wild-type SREBP2 degradation) failed to mediate the degradation of full-length S432A/S436A SREBP2 in functional extracts, indicating that CPD phosphorylation is essential for APC-driven turnover (Fig. 5b); concordantly, FBXW7 depletion (by siRNA) impaired SREBP2 degradation mediated by APC (Fig. 5c, Extended Data. Fig. 5a&b). Together, these data strongly suggest that phosphorylation of SREBP2 within the CPD generates the FBXW7 recognition signal, triggering APC-dependent, FBXW7-mediated ubiquitination and proteolysis of SREBP2. Whether APC directly mediates or facilitates this phosphorylation remains unclear.

**Figure 5.**
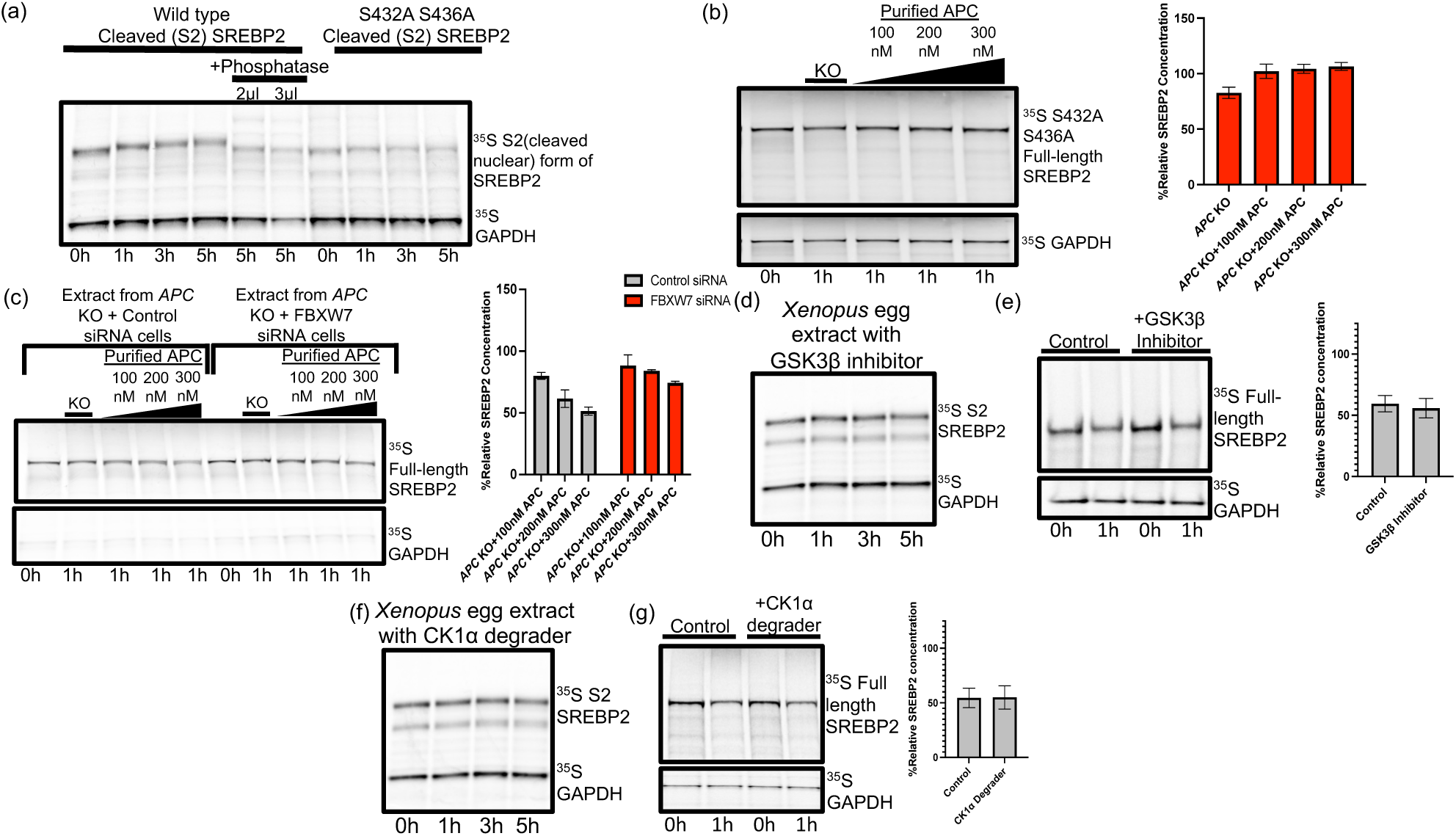
APC promotes SREBP2 degradation via a conserved phosphodegron that licenses recognition by the E3 ubiquitin ligase FBXW7. **a**, The cleaved (S2) form of SREBP2 is phosphorylated in *Xenopus* egg extracts, observed as a progressive mobility shift that is reversed by phosphatase treatment. A S432A/S436A mutant is resistant to this phosphorylation. **b,** Phosphorylation at S432/S436 is required for degradation. In an in vitro degradation assay, full-length S432A/S436A mutant SREBP2 is stable even when *APC* KO lysate is supplemented with high concentrations of purified APC. The bar graph shows quantification from n=3 experiments. No significant differences across 100–300 nM APC (F(2,6)=1.37, p=0.32). **c,** The E3 ligase FBXW7 mediates SREBP2 degradation. siRNA-mediated knockdown of FBXW7 in *APC* KO cells significantly impairs the ability of reconstituted APC to degrade SREBP2. The bar graph shows quantification. Two-way ANOVA showed significant effects of siRNA (F(1,12)=57.7, p<0.00001), APC dose (F(2,12)=27.4, p<0.0001), and their interaction (F(2,12)=4.18, p<0.05), indicating that FBXW7 knockdown blunts APC-dependent degradation. **d-e,** The canonical destruction complex kinase GSK3β is not required. Inhibition of GSK3β with LY2090314 fails to block SREBP2 phosphorylation in *Xenopus* egg extracts (**d**) or its degradation in cell lysates (**e**). Bar graphs show quantification (mean ± s.d.). **f-g,** The canonical kinase CK1α is not required for SREBP2 phosphorylation or degradation. Treatment with a CK1α degrader (SJ3149) fails to block SREBP2 phosphorylation (**f**) or degradation (**g**). Bar graphs show quantification (*mean ± s.d.*).

Strikingly, and in contrast to β-catenin, SREBP2 phosphorylation (S2) and degradation (full-length) did not utilize the canonical destruction-complex kinases, GSK3β or CK1α (Fig. 5d-g, Extended Data Fig. 5c). This differs from SREBP1, whose turnover depends on GSK3β-driven CPD phosphorylation (Extended Data Fig. 5d) (Sundqvist et al., 2005).

### SREBP2 degradation requires the full mutational cluster region of human APC in colon cancers

In humans, APC truncations are commonly found in colorectal cancer cluster within the mutational cluster region (MCR), producing C-terminally shortened proteins (Fig. 6a) (Miyoshi et al., 1992; Fearnhead et al., 2001). To define the APC structural elements required for SREBP2 turnover, we compared full-length APC with three cancer-relevant truncations: T1556* (COLO-205), Q1338* (SW480), and S811* (COLO-320DM) (Extended Data Fig. 6). SREBP2 degradation was abrogated across all C-terminal truncations, indicating a strict requirement for an intact MCR (Fig. 6b, c). By contrast, β-catenin is still degraded by Q1338* APC, albeit with reduced efficiency relative to full-length APC; β-catenin degradation was ultimately lost with the most proximal S811* truncation (Fig. 6d, e). These findings reveal a key mechanistic distinction in APC-dependent degradation of SREBP2 versus β-catenin: SREBP2 turnover requires the entire MCR, whereas β-catenin degradation tolerates substantial but not complete C-terminal loss of this region, as seen in human tumors.

**Figure 6.**
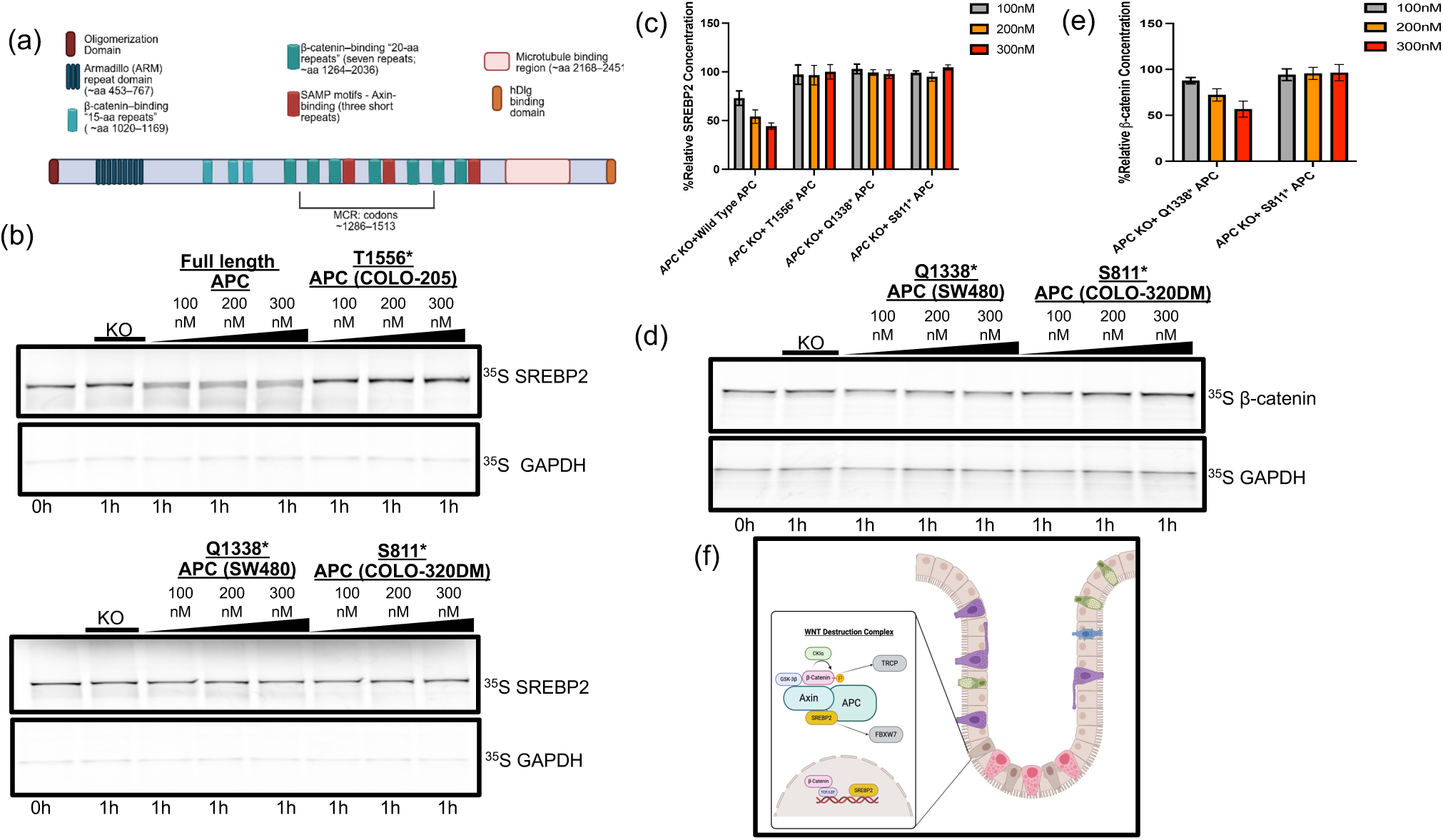
Cancer-associated APC truncations selectively abrogate SREBP2 degradation while retaining activity towards β-catenin. **a**, Schematic of the human APC protein, highlighting key functional domains and the location of the mutation cluster region (MCR), which is frequently deleted in colorectal cancer. **b,** Representative in vitro degradation assays of ³⁵S-labeled SREBP2. Lysates from *APC* knockout (KO) cells were supplemented with increasing concentrations of purified full-length (WT) APC or the indicated C-terminally truncated mutants found in cancer cell lines. Only full-length APC mediates SREBP2 degradation. **c,** Quantification of SREBP2 levels from (b) at the 1-hour time point. Data are *mean ± s.d.*. One-way ANOVA showed a highly significant effect of APC variant on SREBP2 abundance (F(3,32)=53.3, p<0.0001), with only wild-type APC reducing SREBP2, while T1556*, Q1338*, and S811* mutants showed no degradation, confirming that MCR-truncations render APC unable to degrade SREBP2. **d,** Control in vitro degradation assays using ³⁵S-labeled β-catenin. The Q1338* APC mutant, which was inactive against SREBP2, retains its ability to mediate dose-dependent degradation of β-catenin. The more severely truncated S811* mutant is inactive. **e,** Quantification of β-catenin at 1 hr. Q1338* retained β-catenin–degrading activity (ANOVA F(2,6)=16.33, p<0.01; slope = −0.155, p<0.001), whereas S811* showed no APC dose effect (F(2,6)=0.07, p>0.9) and no trend (slope = 0.011, p>0.7). Q1338* differed significantly from S811* (72.3 ± 14.6 vs. 95.4 ± 6.4; p<0.005). Data are *mean ± s.d.*. **f,** Role of the Wnt destruction complex in intestinal epithelial cells. A magnified view highlights the molecular components of the WNT destruction complex, targeting β-catenin and SREBP2 for degradation. These degradation processes help regulate Wnt signaling and cholesterol metabolism, maintaining homeostasis in the intestinal epithelium.

## DISCUSSION

β-Catenin is the only well-validated direct substrate of the Wnt destruction complex whose stability is regulated by extracellular Wnt ligands. Here, we challenge this β-catenin-centric view by identifying a novel, yet important, target of Wnt pathway regulation, SREBP2—the well-studied master regulator of cholesterol metabolism. We show that SREBP2 is a direct, β-catenin independent, substrate of the WNT destruction complex. Using a reconstituted cell-free system and in vivo models from *Xenopus* embryos, mouse colon, and human colorectal cancer cell lines, we show that the tumor suppressor APC and its partner AXIN1 promote SREBP2 degradation. This discovery expands the complex’s substrate repertoire. Furthermore, it forges a direct mechanistic link between developmental signaling, cancer biology, and cellular metabolism (Fig. 6f).

Mechanistically, SREBP2 regulation diverges from β-catenin. Both substrates engage the APC–AXIN1 scaffold, but their degradation is licensed by distinct phosphorylation events: β-catenin requires CK1α and GSK3β, whereas SREBP2 depends on phosphorylation that creates an FBXW7-recognized phosphodegron and is independent of CK1α/GSK3β. Furthermore, unlike SREBP2, but similar to β-catenin, SREBP1 requires GSK3β mediated phosphorylation to be degraded by FBXW7 (Sundqvist et al., 2005), likely reflecting differences in the amino acid sequence surrounding their phosphodegrons (SREBP1’s T426/S430 versus SREBP2’s S432/S436). These findings also suggest how cells can control cholesterol or fatty acid pathways in different physiological contexts.

The discovery that APC and AXIN1 co-regulate both SREBP2 and β-catenin reframes each as master signal integrators, not just downstream components of the same linear pathway. By acting as molecular scaffolds, APC and AXIN1 can simultaneously control distinct downstream effector circuits. This allows them to translate a common upstream Wnt signal into multifaceted cellular responses. The ramifying Wnt signal with some differential posttranslational controls may directly control several regulatory processes. Such integration at the posttranscriptional and posttranslational level may be essential for complex processes like development and stem cell regulation, which might demand the coordinated activation of both transcriptional programs (governed by β-catenin) and the metabolic machinery (governed by SREBP2). This raises an intriguing possibility that the WNT mediated destruction complex may recruit an even wider array of substrates to orchestrate other key physiological processes and that APC and AXIN1 scaffolds may assemble an array of substrate-specific subcomplexes to orchestrate other key physiological processes, all coordinated by extracellular WNT signaling. We have some preliminary evidence that this is true.

It was most intriguing to find that cancer-relevant truncations that delete the entire distal mutational cluster region (MCR) completely abrogate SREBP2 degradation while still retaining some capacity to degrade β-catenin. This dichotomy offers a powerful explanation for the predominance of certain APC mutations, such as T1556* and Q1338*, in colorectal cancer that leave β catenin subject to degradation but block SREBP2 degradation. This argues that tumor selection is not merely for β-catenin stabilization, but also for the complete loss of APC’s metabolic surveillance—a function exquisitely sensitive to MCR truncation.

The metabolic consequences of this escape from APC-mediated control may provide a direct link to oncogenic signaling. In an APC-deficient colonic epithelium, *de novo* cholesterol synthesis is significantly elevated (Muta et al. 2023), and prior studies confirm that enhanced cholesterol production through SREBP2 copy-number gain is a potent driver of intestinal tumorigenesis in *Apc^min^* mice (Wang et al. 2018). It should be noted that increased flux through the mevalonate pathway also elevates levels of the isoprenoid geranylgeranyl pyrophosphate (GGPP), which is required for the prenylation and membrane localization of small GTPases like KRAS (Muta et al. 2023). In colorectal tumors, where *APC* loss frequently co-occurs with activating KRAS mutations (Lee et al. 2022), this GGPP overexpression would be poised to hyperactivate the MEK/ERK cascade. This rewiring could create a potent feed-forward circuit: truncated APC disables SREBP2 control, elevating GGPP and amplifying RAS–ERK signaling, which could be a mechanistic rationale for the strong selection for MCR-truncating mutations in colorectal tumors.

Beyond its role in intestinal cancer, this work invites exploration into the broader physiological consequences of Wnt-dependent metabolic control (Sethi & Vidal-Puig, 2010). The APC-SREBP2 axis provides a mechanism to couple proliferative cues with the anabolic capacity to build new tissues—a link especially critical in lipid-rich environments like the developing nervous system, where cholesterol availability governs progenitor proliferation, synapse formation, and myelination (Roy & Tedeschi, 2021).

Future work using in vivo genetic models will be crucial for answering key physiological questions, such as how this axis contributes to the maintenance of adult stem cell niches, where tissue stem cells must balance metabolic readiness with proliferative restraint. Further these recent studies suggest a potential metabolic arm to cancer induction and maintenance. Ultimately, defining the role of this newly uncovered axis in both normal physiology and cancer promises to expose a new set of pharmacologic and dietary vulnerabilities for therapeutic intervention.

### METHODOLOGY

#### Cell Lysate Preparation

Cells were grown in 20-cm dishes to 80–90% confluence. Dishes were placed on ice and washed three times with 10 ml ice-cold hypotonic buffer (25 mM Tris-HCl pH 7.5, 10 mM KCl, 1.5 mM MgCl₂). After the final wash, residual buffer was aspirated completely. Cells were scraped into 15-ml conical tubes and pelleted (4,000 × g, 5 min, 4°C). Pellets were resuspended in digitonin lysis buffer at 0.75 volumes relative to packed cell volume (25 mM Tris-HCl pH 7.5, 50 mM NaCl, 10 mM MgCl₂, 4 mM ATP, 1 mM TCEP, 15% glycerol, 0.1% [w/v] digitonin) supplemented with protease and phosphatase inhibitors. Lysates were clarified (4,000 × g, 5 min, 4°C) to remove nuclei and insoluble material. The supernatant (nucleus-free cytosolic extract) was collected, protein concentration determined by 660nm Protein Assay Reagent (Thermo Fisher), aliquoted, flash-frozen in liquid nitrogen and stored at −80 °C (single freeze–thaw).

#### In vitro degradation assays

Reactions (20 µl total) contained 14 µl cytosolic extract, 2 µl ATP-regenerating mix (final: 7.5 mM creatine phosphate, 1 mM ATP, 0.1 mM EGTA, 1 mM MgCl₂, pH 7.7), 1 µl ubiquitin (from a 50 mg ml⁻¹ stock), and 1 µl ^35^S-methionine-labelled substrate generated with the TNT SP6 Quick Coupled Transcription/Translation System (Promega). Where indicated, recombinant APC and/or Axin1 were added at the specified concentrations.

Reactions were incubated at 30°C for the indicated times and quenched with 4× LDS sample buffer (Genscript, M00676-250). Samples were resolved by SDS–PAGE, gels were dried and exposed to a phosphor screen, and signals were visualised by autoradiography and quantified in ImageJ relative to t₀.

#### Pharmacological perturbations

For kinase inhibition, Expi293 cells were pre-treated for 4 h with LY2090314 (GSK3β inhibitor, 2 µM) (Atkinson et al. 2015) or SJ3149 (CK1α degrader, 2 µM) (Nishiguchi, G. et al. 2024). For proteostasis controls, cells were treated with carfilzomib (proteasome inhibitor, 10 µM, 4 h) (Kim & Crews 2013) or TAK-243 (E1 inhibitor, 5 µM, 1 h; shortened to limit proteotoxic stress) (Hyer et. al. 2018). Where indicated, UBOK (E1 inhibitor) was added directly to reactions to a final concentration of 100–500 µM. For *Xenopus* lysates, all inhibitors were used at 10 µM.

#### Cell culture and sterol pre-conditioning

Control and isogenic APC-KO cells were maintained in DMEM supplemented with 10% FBS, 1% penicillin–streptomycin, at 37°C/5% CO₂. To deplete intracellular cholesterol, cells were cultured overnight (∼16 hours) in a base medium appropriate for the cell line, supplemented with 5% lipoprotein-deficient fetal bovine serum (LPDS; Biowest, S148L), 5 µM simvastatin (AdipoGen, AG-CN2-0539), and 10 µM mevalonate (Millipore-Sigma, M4667). Before adding the deprivation medium, cells grown to 70–80% confluence were washed once with sterile PBS to remove residual serum.

For WNT stimulation, cells were sterol-deprived for 16 h, then treated with FZD7/8-subtype NGS WNT (Kaktus Bio, WNT-HM23A) and RSPO1 (Sino Biological, 11083-H08H) at 50 ng/mL each.

### Subcellular Fractionation

Cells were harvested, washed with PBS, and resuspended in hypotonic lysis buffer (50 mM Tris HCl, pH 7.8; 1 mM EDTA with fresh protease inhibitors). For lysis, the cell suspension was subjected to nitrogen cavitation using a Parr cell disruption vessel.

Cells were equilibrated with nitrogen gas at 800 psi for 20 minutes at 4°C, followed by rapid decompression to lyse the cells while leaving nuclei intact. Nuclei were then pelleted by centrifugation at 1,000 x g for 5 minutes. The resulting supernatant was subjected to ultracentrifugation at 100,000 x g for 1 hour to pellet the membrane fraction, which was designated the Membrane Fraction. The nuclear pellet was resuspended in a high-salt nuclear extraction buffer (20 mM HEPES/KOH, pH 7.6; 2.5% glycerol; 1.5 mM MgCl₂; 0.42 M NaCl; 1 mM EDTA; 1 mM EGTA) and rotated for 1 hour at 4°C. After centrifugation, the supernatant was collected as the Nuclear Extract. Protein concentrations were determined for both fractions prior to immunoblotting.

#### Immunoblotting

Equal amounts of protein were separated on SurePAGE™, Bis-Tris, 10×8, 4-12%, 12 wells (GenScript, M00653) and transferred to PVDF membrane (Millipore Sigma, Immobilon-FL, IPFL85R). Membranes were blocked for 1 hour with Intercept (PBS) Blocking Buffer and incubated overnight at 4°C with primary antibodies dissolved in Intercept (PBS) Blocking Buffer with 0.2% Tween 20. After washing, membranes were incubated with species-specific, fluorophore-conjugated secondary antibodies (1:10,000 dilution) for 1 hour at room temperature. Blots were imaged using a fluorescence imaging system (LI-COR Odyssey CLx) and quantified with Image Studio Lite software. β-Actin served as the loading control.

Following primary antibodies were used: mouse anti-APC (1:1000, Usbio, A2298-70A), goat anti-SREBP2 (1:2000, R&D Systems, AF7119), rabbit anti-SREBP1 (1:2000, Proteintech, 14088-1-AP), rabbit anti-Lamin A/C (1:5000, Proteintech, 10298-1-AP), mouse anti-Calnexin (1:2000, Proteintech, 66315-1-Ig), Rabbit anti-LDLR (1:2000, Proteintech, 10785-1-AP), rabbit anti-HMGCR (1:1000, Proteintech, 13533-1-AP), rabbit anti-FBXW7 (1:1000, Proteintech, 28545-1-AP), rabbit anti-FDPS (1:1000, Proteintech, 16129-1-AP), rabbit anti-FDFT1 (1:1000, Proteintech, 13128-1-AP), rabbit anti-HMGCS1 (1:1000, Proteintech, 17643-1-AP), rabbit anti-MVK (1:1000, Proteintech, 12228-1-AP), mouse anti-β-catenin (1:2500, BD Biosciences, 610154), and mouse anti-β-actin (1:5000, Proteintech, CL750-81115).

#### Xenopus Embryo Microinjection and Analysis

*Xenopus* laevis eggs were obtained, fertilized in vitro, and de-jellied in 2% cysteine. Purified, Strep II-tagged human APC protein was injected at a final concentration of 10, 20, and 30nM in a buffer containing Texas Red dextran as a tracer.

Single-cell stage embryos were microinjected with 15nL of the APC solutions or a buffer-only control. At stage 6.5, groups of 10 embryos were snap frozen. The embryos were then lysed by Dounce homogenization in a hypotonic buffer, and the cytoplasmic fraction was isolated by centrifugation.

Protein lysates were normalized for total protein content and analyzed by immunoblotting. Proteins were separated on 4–12% Bis-Tris gels, transferred to PVDF membranes, and probed with antibodies against SREBP2 and β-actin.

#### RNA Extraction and RT–qPCR

Total RNA was extracted by a TRIzol-based phase-separation method (Thermo Fisher Scientific). Contaminating genomic DNA was removed using the TURBO DNA-free™ Kit (Invitrogen).

RT-qPCR was performed with the LunaScript Multiplex One-Step RT-PCR Kit (NEB). Reactions were run with the following thermal cycling conditions: 55°C for 10 min, 95°C for 3 min, followed by 40 cycles of 95°C for 10 s and 60°C for 30 s. Relative expression was calculated using the ΔΔC_t_ method, with normalization to Actb/ACTB. All results are from at least three biological replicates.

Primer sequences (5′→3′, F/R).

Mouse: Actb F GATTACTGCTCTGGCTCCTAG, R GACTCATCGTACTCCTGCTTG; FDFT1 F ACTTCCACACTTTCCTCTATGAC, R CTCAGCCAAATTTCTAAACTCCAG; FDPS F GGAGCCGAAGAAACAGGATG, R CTGACACAAGGAAGAAAGCCT; HMGCR F CACATTCACTCTTGACGCTCT, R ACTGACATGCAGCCGAAG; LDLR F AGACCCAGAGCCATCGTAG, R GATGTCCACACCATTCAAACC; MVK F GTGAGCGTCAATTTACCCAAC, R TACCGAGACATCACCTTGC; HMGCS1 F CTGCTATTCTGTCTACCGCAA, R TGAGTGAAAGATCATGAAGCCA.

Human: ACTB F ACAGAGCCTCGCCTTTG, R CCTTGCACATGCCGGAG; FDFT1 F ACAACCTGGTGCGCTTC, R GATAACAGCTGCGAAACTGC; FDPS F CAGCTTTCTACTCCTTCTACCTT, R CTGAAAGAACTCCCCCATCTC; HMGCR F GTTTACCCTCGATGCTCTTGT, R CTGACATGCAGCCAAAGC; LDLR F TGGCAGAGGAAATGAGAAGAAG, R AAAGTTGATGCTGTTGATGTTCT; MVK F CTGCTCAAGTTCCCAGAGAT, R TCATGTCAATGAGCTCTTCCA; HMGCS1 F GATGAAGGAGTAGGACTTGTGC, R CCTCACAGAGTATCTTAATGTTCCC.

#### Mouse strains

Mice were under the husbandary care of the Department of Comparatie Medicine at the Koch Institute for Integrative Cancer Research. All mice experiments were approved by the Massachusetts Institute of Technology (MIT) Committee on Animal Care and conducted in accordance with institutional guidelines.

The *Apc^flox/flox^;Villin^CreERT2^* mice used in this study were generated by crossing *Apc^flox/flox^* mice to *Villin-CreERT2* mice. This model allows for tamoxifen-inducible deletion of *Apc* specifically in the intestinal epithelium, driven by the Villin promoter. The *Apc^flox/flox^* allele contains loxP sites flanking exon 14 (Colnot et. al. 2004). In this study, both male and female mice were used at 2–4 months of age unless otherwise specified in the figure legends.

#### Colonoscopy and Focal Gene Deletion

For focal gene deletion in the distal colon, mice were anaesthetized with isoflurane and imaged using a high-resolution mouse video colonoscope (Karl Storz). The lumen was gently flushed with sterile PBS to clear fecal material, after which the endoscope was advanced ∼1–2 cm. A 30-gauge needle passed through the instrument channel delivered 50 µL 4-hydroxy tamoxifen (100 uM) into the submucosa, producing a visible bleb that confirmed deposition. This single submucosal injection reliably induced Cre-mediated recombination in the overlying epithelium, generating a localized *Apc*-deficient patch; the untreated proximal colon of the same animal served as a matched isogenic control. Mice were monitored continuously until full recovery from anasthesia.

At 2 weeks post-injection, distal lesional tissue and matched proximal controls were collected for RT–qPCR, immunofluorescence, and Western blot analyses.

#### Immunoflorescense

Following euthanasia, mice were perfused with ice-cold PBS + heparin (10 U mL⁻¹); colons were resected, cut into 4–5 mm pieces, fixed in 4% formaldehyde (24 h, 4°C), rinsed ×3 in cold PBS, and stored in PBS + 0.02% sodium azide (4 °C). Tissues were embedded in 2.5% agarose and sectioned (Compresstome ZF-310-0F) into 120 µm slices (speed 2; oscillation 6). Slices were optionally pre-permeabilized in 0.2–0.3% Triton X-100 (10–15 min, RT), then permeabilized in 0.1% Tween-20 (30–45 min) and blocked overnight at 4 °C in PBS + 5% normal donkey serum + 0.05–0.1% Tween-20 + 0.02% azide. Primary antibodies were applied overnight at 4°C in PBS + 10 µM CaCl₂ + 0.02% azide + 0.05–0.1% Tween-20: rabbit anti-β-catenin (Proteintech 51067-2-AP; 1:700) and goat anti-mouse SREBP2 (R&D AF7119; 1:1000). After washes (6 × 30 min, cold PBS + 0.02% azide + 0.05–0.1% Tween-20), secondaries (donkey anti-rabbit AF594; donkey anti-goat DL405) were incubated overnight at 4°C in the same buffer, followed by identical washes. Nuclei were stained with TO-PRO-3 overnight at 4°C in PBS + 10 µM CaCl₂ + 0.02% azide, washed (6 × 30 min, PBS), and mounted in antifade medium (0.2% propyl gallate in PBS with 0.8% d₆-DMSO and 10 µM CaCl₂). Imaging was on an Olympus FV3000 (60×/1.2 NA water; ∼414 nm pixel size).

#### Colonic Crypt Isolation

Colonic crypts were isolated from control (proximal) and *Apc*-deficient (distal) regions for downstream analysis. Following euthanasia, the colon was dissected, and the proximal and distal segments were separated and processed independently. Each tissue segment was opened longitudinally, washed with cold PBS, and cut into ∼ 1 cm pieces. The pieces were then incubated in PBS with 10 mM EDTA for 15 minutes at 37°C on a shaker to release the epithelial fraction. After incubation, tissue was transferred to fresh PBS, and crypts were dislodged by vigorous shaking. The resulting suspension was pelleted, washed once, and were immediately used for cholesterol isolation.

#### Cholesterol extraction and quantification

Before extraction, a small aliquot of each crypt suspension (10–20 µL) was reserved for protein quantification; the remainder was kept on ice. Intestinal crypts in PBS were pelleted (500 x g, 5 min, 4°C), supernatant removed, and the wet pellet (≤0.15 g) extracted per instructions (Sigma-Aldrich, MAK175): 3 mL Extraction Solvent, homogenize/vortex, add 0.5 mL Aqueous Buffer, then pass through the kit syringe filter to collect the eluate. Eluates were dried under N₂ (30–37 °C) and re-solubilized in Amplex Red reaction buffer (Thermo Fisher, A12216). Cholesterol content was measured with Amplex Red Cholesterol Assay kit (Thermo Fisher scientific, #A12216).

#### [^14^C] acetate sterol synthesis assay

*De novo* sterol synthesis was measured by pulsing cells at ∼70–80% confluence—after overnight sterol depletion—with sodium [1-^14^C]acetate (2.5 µCi mL⁻¹; final ethanol ≤0.25% v/v) for 2 h at 37 °C. After three ice-cold PBS washes, cells were pelleted (500 × g, 5 min, 4°C) and lysed; a 10% aliquot was reserved for protein quantification at 660 nm, and the remaining lysate was normalized accordingly before saponification in 1 M KOH/90% ethanol with 0.01% BHT at 80 °C for 1–2 h. Neutral sterols were extracted with 99% hexane (0.01% BHT), dried under nitrogen, and re-dissolved for TLC (hexane: diethyl ether: acetic acid, 90:10:1). Radiolabeled cholesterol was visualized by phosphor imaging (24–72 h) alongside cold standards and quantified in ImageQuant/ImageJ.

#### siRNA-mediated Knockdown

For knockdown of FBXW7, cells were transfected with a pool of three distinct siRNA duplexes (Integrated DNA Technologies) at a final concentration of 20 nM. Transfection was performed using Lipofectamine RNAiMAX (Thermo Fisher Scientific) according to the manufacturer’s protocol. A non-targeting scrambled siRNA was used as a negative control. Cells were harvested for analysis or subsequent experiments 48 hours post-transfection. Knockdown efficiency was confirmed by Western blot.

siRNA Sequences for FBXW7 knockdown hs.Ri.FBXW7.13.1**

Sense:** ‘5’-rArUrCrUrGrCrArArGrCrArArArArGrCrUrCrArUrUrUrUrUCT-3’’ Antisense:** ‘5’-rArGrArArArArUrGrArGrCrUrUrUrUrUrGrCrUrUrGrCrArGrArUrC-3’’ hs.Ri.FBXW7.13.2**

Sense:** ‘5’-rGrUrGrArGrCrCrGrUrUrUrArCrArGrGrUrUrC rArC rArAGA-3’’

Antisense:** ‘5’-rU rCrU rUrGrUrG rArG rArArC rUrGrU rArArArC rGrGrC rUrCrArC rArG-3’’hs.Ri.FBXW7.13.3**

Sense:** ‘5’-rArArCrArUrUrGrCrArArGrGrUrCrCrArArCrArArGrCAT-3’’ Antisense:** ‘5’-rArUrGrCrUrUrGrUrUrGrGrArCrCrUrUrGrCrArArUrGrUrUrG-3’’

#### Statistics and reproducibility

Unless stated otherwise in the main text or figure legends, all experiments were repeated independently at least three times, and *n* refers to the number of biological replicates. No samples, data points, or animals were excluded from analysis, and formal a priori sample size calculations were not performed. Age- and sex-matched mice were randomly assigned to experimental groups. Experiments were generally not performed under blinded conditions. Data are presented as mean ± s.d. unless indicated otherwise. Comparisons between two groups were carried out using unpaired, two-tailed Student’s *t*-tests; where matched samples were analysed, paired two-tailed *t*-tests were used as specified. For experiments involving more than two groups or multiple factors, one-way or two-way ANOVA and linear regression were applied as indicated in the figure legends. All statistical analyses were conducted using GraphPad Prism 9. Exact *n* values, statistical tests, and *p* values are reported in the figure legends.

## ACKNOWLEGEMENT

We thank Hong Kang for help in organizing the human transcription factor library for screening, Bai Luan for the pSNAFf 1×FLAG–AXIN1–SNAP–HALO–His plasmid and guidance on Axin1 purification, and William Ratzan and Christine Field for help with *Xenopus* microinjections. We thank Michael Ranes and Sebastian Guettler for pLIB_dStrepII-TEV-APC construct. We are grateful to Peter Tontonoz and John Paul Kennelly for advice and for sharing the sterol-deprivation protocol, and to Ying Lu, Ramesh Shivdasani, Bruce Spiegelman, and Rajat Rohatgi for valuable insights and suggestions. We thank Initiative for Genome Editing and Neurodegeneration for generating CRISPR edited cell lines. A.R. was supported by the DF/HCC SPORE in Gastrointestinal Cancer Career Enhancement Program (NIH P50 CA127003). Ö.H.Y. is supported by the US National Institutes of Health (R01CA245314, R01CA257523, R01DK133919, R01DK140310, P30CA014051, R01DK126545, and 3OT2CA297570), the Kathy and Curt Marble cancer research award, a Koch Institute–Dana-Farber/Harvard Cancer Center Bridge project grant and AFAR; and receives support from the MIT Stem Cell Initiative. M.W.K. is supported by an NIH MIRA (R35 GM145248).

## AUTHORS CONTRIBUTION

Conceptualization: A.R., M.K.; in vitro degradation assay development: A.R.; investigation: A.R., C.R., W.T., and E.G.; mouse models and colonoscopy induced gene deletion: S.I., S.K., H.S., and O.H.Y.; Xenopus experiments: C.A.; analysis and interpretation of data: A.R., C.R., O.H.Y., M.K.; writing – original draft: A.R.; writing – review and editing: M.K., and O.H.Y., and other authors.

