## Supplementary figures and data for "The Wnt/APC destruction complex targets SREBP2 in a β-catenin-independent pathway to control cholesterol metabolism"

**Extended Data Figure 1**

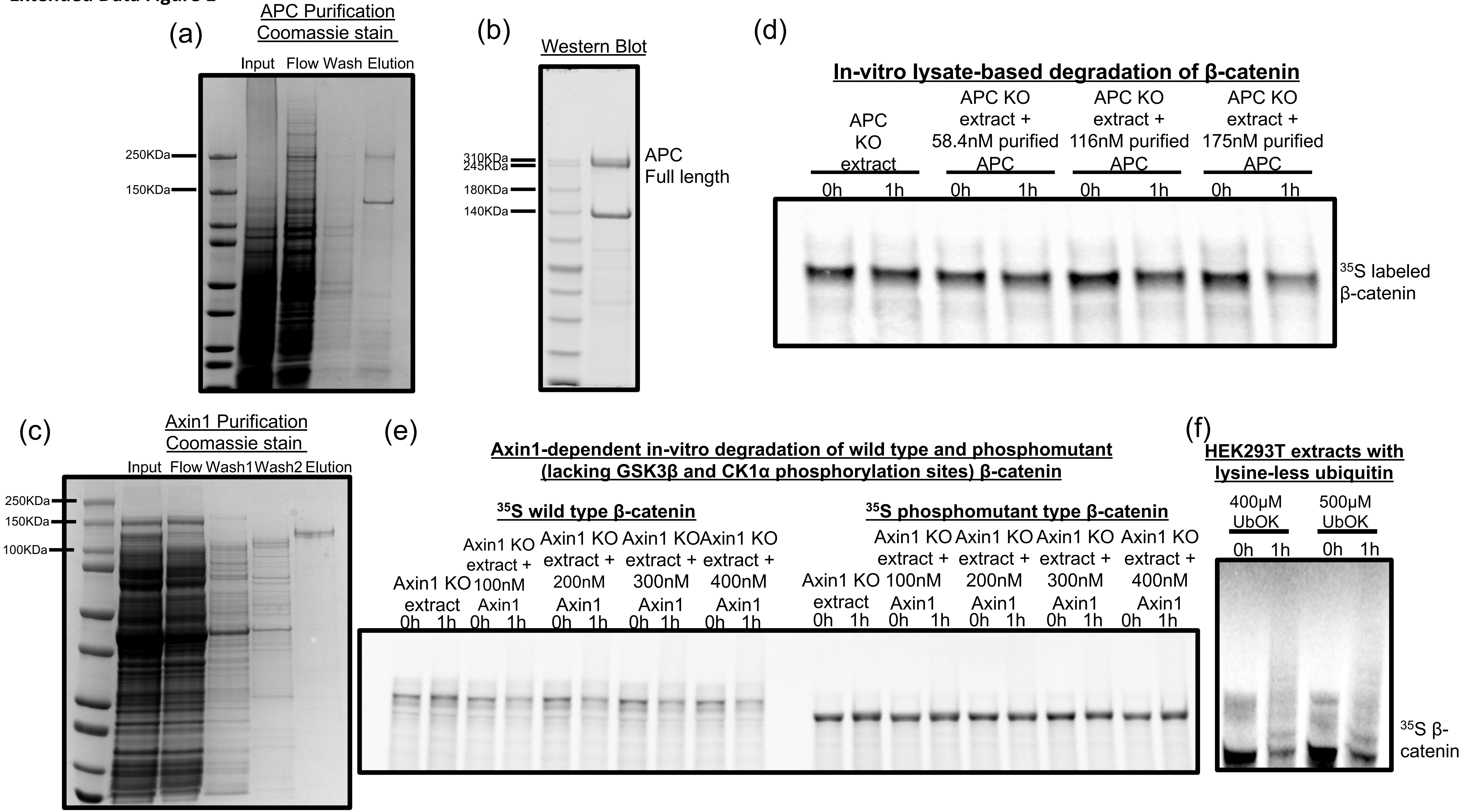

Extended Data Figure 2

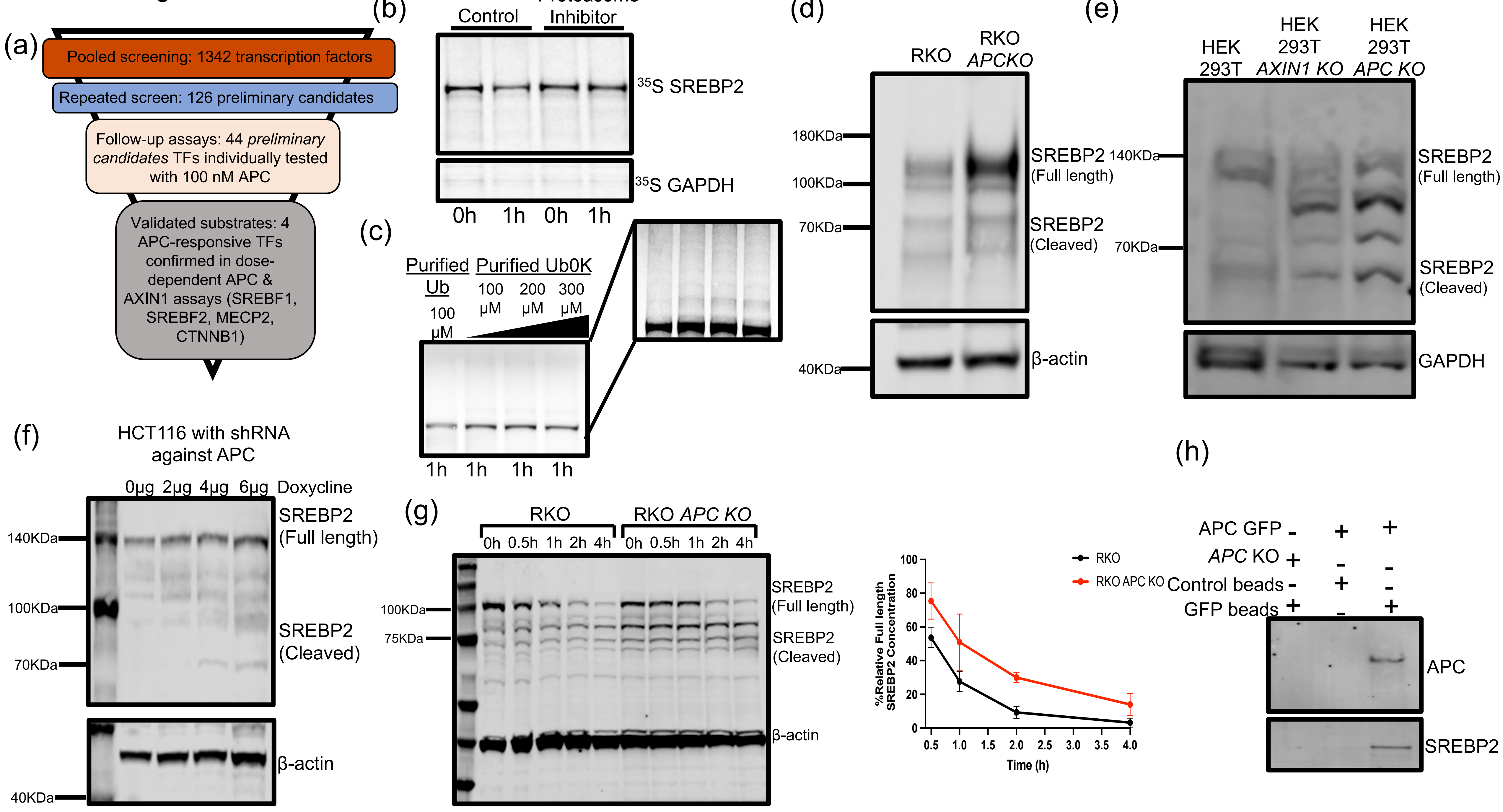

**Extended Data Figure 3**

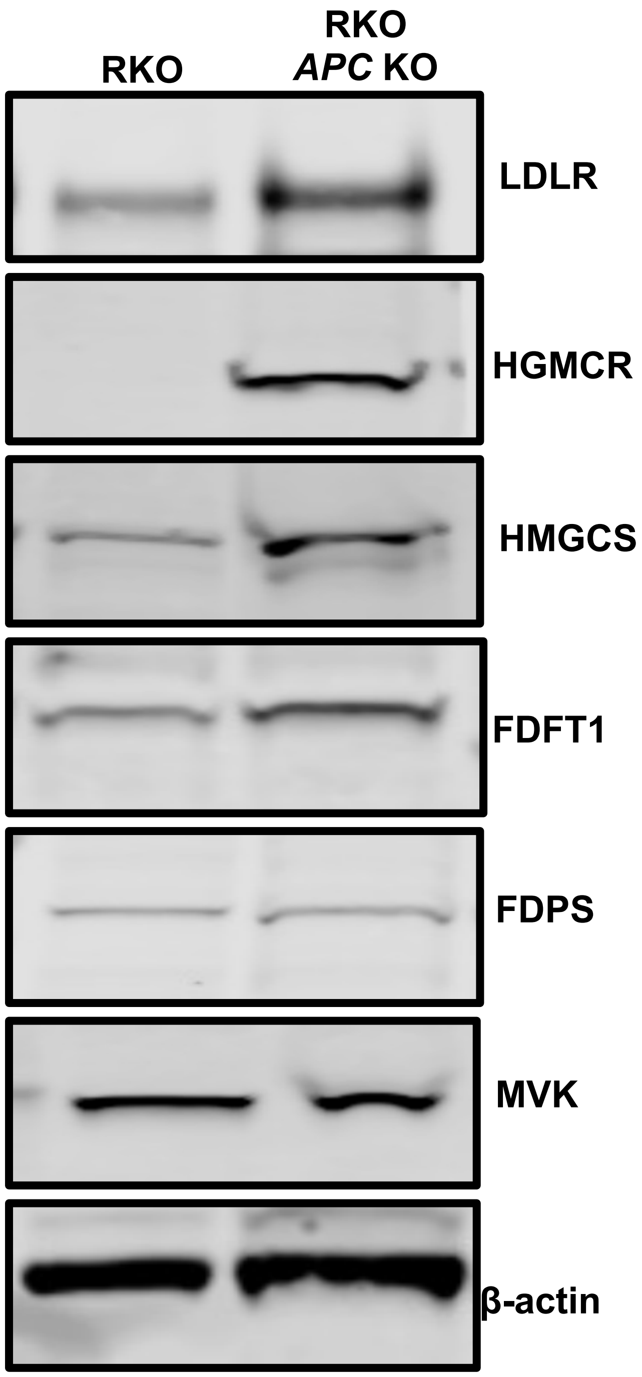

**Extended Data Figure 4**

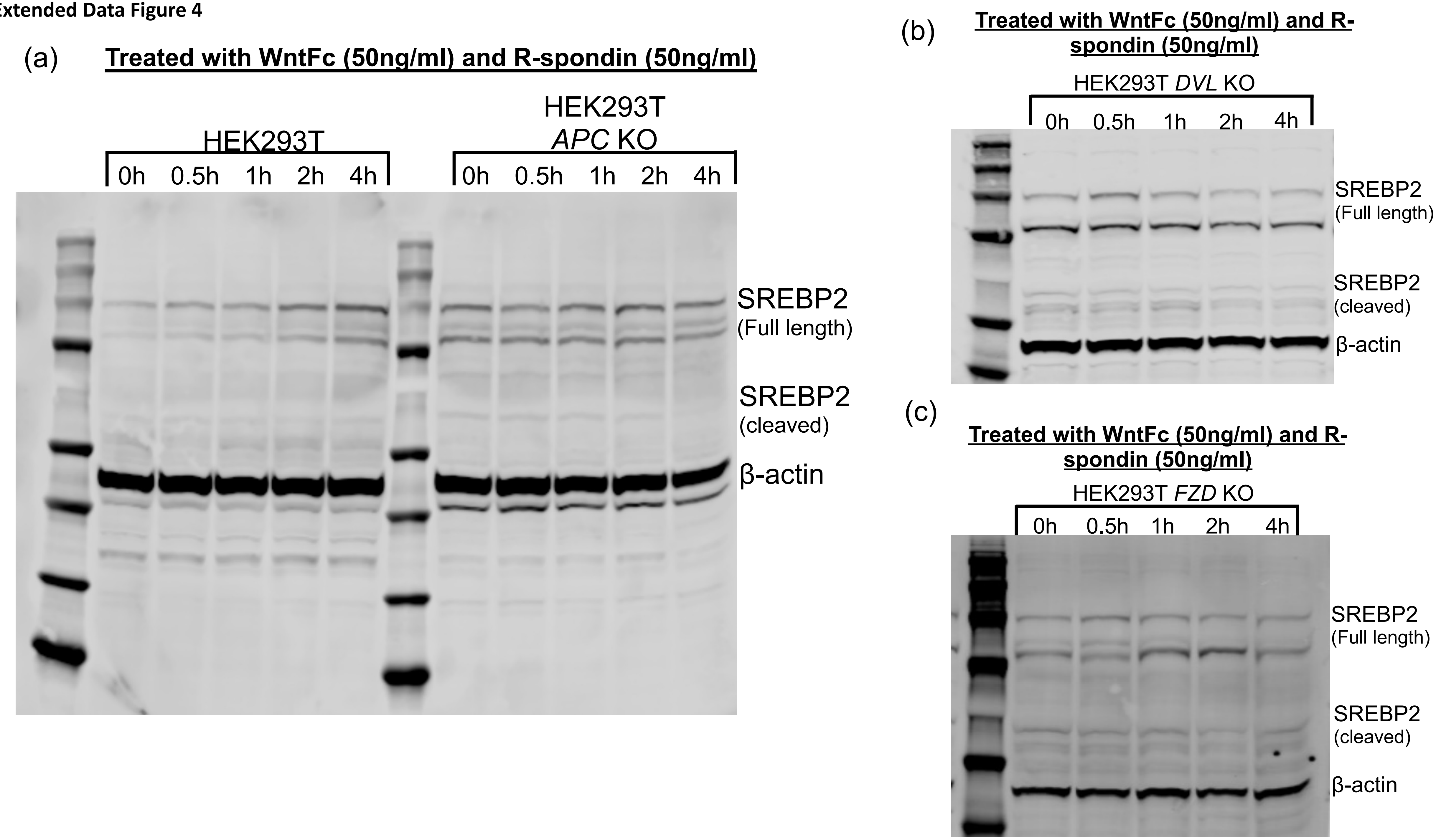

**Extended Data Figure 5**

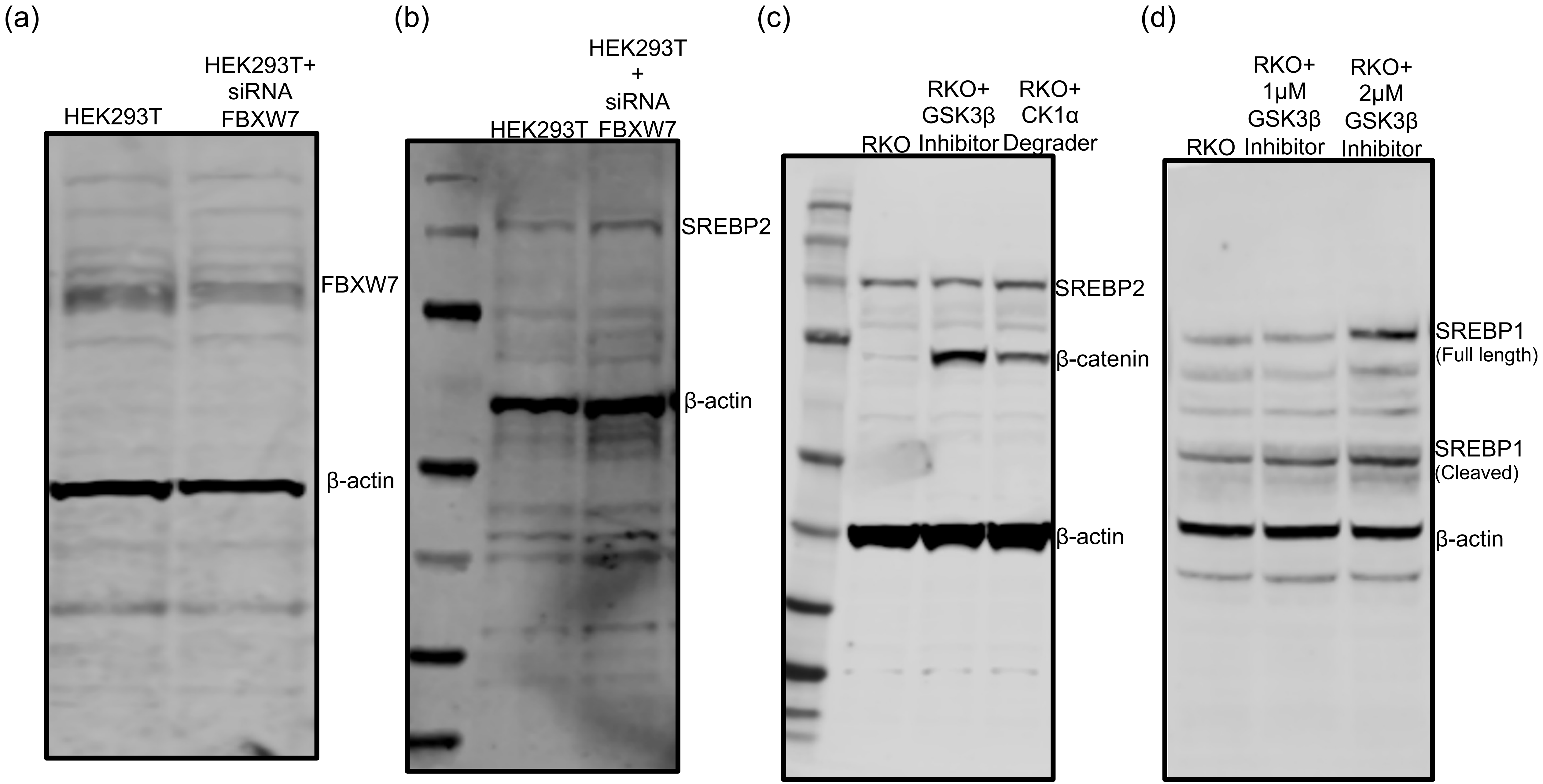

**Extended Data Figure 6**

Purified C-terminus truncated APC

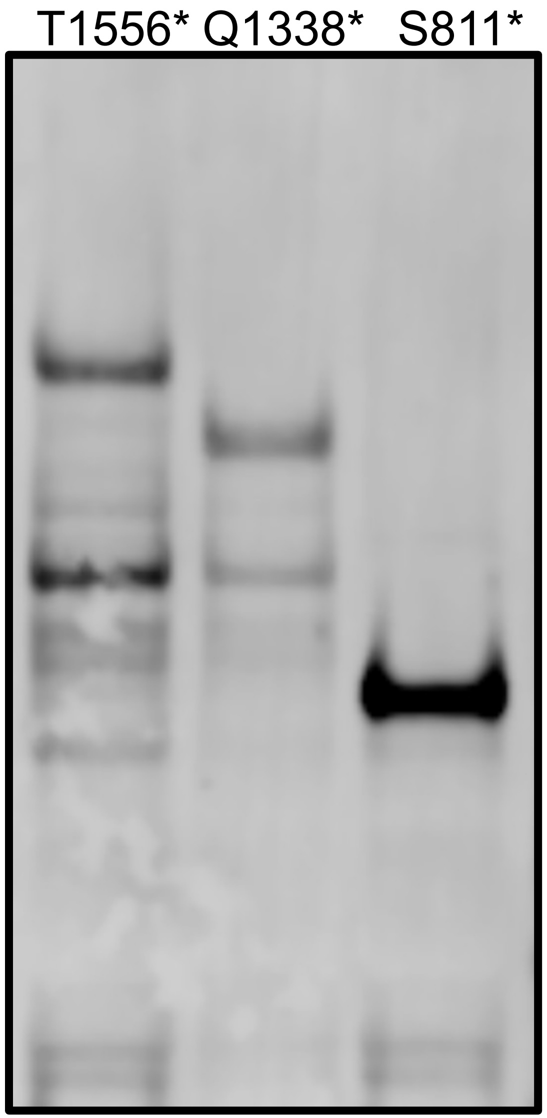

### Extended Data Method

#### Recombinant APC expression and purification

Full-length human APC containing an N-terminal StrepII tag (pLIB-StrepII-APC) was kindly provided by Sebastian Guettler.[1] Recombinant baculoviral bacmids were generated by Tn7 transposition in *DH10 MultiBacTurbo E. coli* competent cells as described for the MultiBac system.[2] Bacmids were purified and used to transfect adherent Sf9 cells ($5 \times 10^5$ cells $mL^{-1}$) grown in ESF 921 Insect Cell Culture Medium, protein-free (Expression Systems, cat. 96-001-01) supplemented with 1% penicillin–streptomycin (Thermo Scientific, cat. 15070063). For initial transfection, 2 mL of cells were transfected with purified bacmid DNA using Cellfectin II (Thermo Scientific, cat. 10362100). P1 virus was harvested after 72 h at 27°C without agitation. The entire ~2 mL of P1 virus-containing medium was used to infect 30 mL of Sf9 cells at $5 \times 10^5$ cells $mL^{-1}$ for a second amplification at 27°C with shaking at 130 rpm. P2 virus was collected once cell viability reached ~80% and stored at 4°C.

For protein production, infected cells were lysed in I-PER™ Insect Cell Protein Extraction Reagent (Thermo Scientific, cat. 89802) supplemented with cOmplete™, Mini, EDTA-free protease inhibitors (Sigma, cat. 11836170001). Lysates were clarified sequentially at 4,000 g (10 min) and 20,000 g (30 min). Cleared supernatants were loaded onto a Strep-Tactin 4Flow high-capacity FPLC column (IBA Lifesciences, cat. 2-1258-001), washed with 30 column volumes of wash buffer (100 mM Tris-Cl, 150 mM NaCl, 10 mM EDTA, pH 8.0), and eluted with 50 mM desthiobiotin. Eluates were dialysed overnight at 4°C (Slide-A-Lyzer™ G3, 2 kDa MWCO, Thermo Scientific, cat. A52962) against 25 mM Tris-HCl pH 7.5, 50 mM NaCl, 10 mM $MgCl_2$, 4 mM ATP, 1 mM TCEP and 15% glycerol. After dialysis, samples were snap-frozen in liquid nitrogen and store at -80°C.

#### AXIN1 expression and purification

HEK293-derived Expi293 cells were transiently transfected with the pSNAFf 1×FLAG–AXIN1–SNAP–HALO–His plasmid. For small-scale optimization (2 mL), 2–3 µg DNA was used (≈1–1.5 µg $mL^{-1}$). For preparative expression in a 200 mL culture at ~$2 \times 10^6$ cells $mL^{-1}$, 200–300 µg plasmid DNA (≈1–1.5 µg $mL^{-1}$) was diluted in 150 mM NaCl (or PBS) and complexed with linear PEI (25 kDa) at a 1:2–1:3 DNA:PEI (w/w) ratio. DNA and PEI solutions were incubated separately for 5 min, combined, incubated 15–20 min at room temperature to allow complex formation, and then added dropwise to the culture. Cells were harvested 48 h post-transfection by centrifugation, washed once with ice-cold hypotonic solution, and resuspended in lysis buffer (25 mM Tris-HCl pH 7.5, 50 mM NaCl, 10 mM $MgCl_2$, 4 mM ATP, 1 mM TCEP, 15% glycerol, 10 mM imidazole) supplemented with protease inhibitors.

Cells were lysed by sonication on ice. Lysates were clarified by centrifugation at 20,000 g for 30 min at 4°C, then loaded onto a His60 Ni Superflow Cartridge (Takara, cat. 635680) equilibrated in lysis buffer (10 mM imidazole). The column was washed with 12 column volumes of wash buffer (25 mM Tris-HCl pH 7.5, 50 mM NaCl, 10 mM MgCl, 35 mM imidazole). Bound protein was eluted in the same buffer containing 250 mM imidazole. Elution fractions were pooled and dialyzed overnight at 4°C against storage buffer identical to the lysis buffer but lacking imidazole (25 mM Tris-HCl pH 7.5, 50 mM NaCl, 10 mM $MgCl_2$, 4 mM ATP, 1 mM TCEP, 15% glycerol). Protein was concentrated as needed, aliquoted, snap-frozen in liquid nitrogen, and stored at −80 °C.

**Cycloheximide pulse–chase to measure SREBP2 stability**

RKO and RKO APC-knockout (KO) cells were seeded 24 h before treatment to reach ~70–80% confluence. To synchronise SREBP2 processing, cells were cultured for 16 h in sterol-depleted medium (DMEM + 10% lipoprotein-deficient serum), then switched to complete growth medium immediately before cycloheximide addition. Cycloheximide (InSolution™ Cycloheximide, MilliporeSigma/Calbiochem, cat. 239765-1ML; CAS 66-81-9) was added to a final concentration of 100 µg/mL (t = 0). Cells were harvested at 0, 0.5, 1, 2, and 4 h.

At each time point, plates were rinsed twice with ice-cold PBS and lysed on ice in RIPA Lysis and Extraction Buffer (Boston BioProducts, cat. BP-115D) supplemented with EDTA-free protease inhibitors. Lysates were briefly sonicated on ice (short bursts to shear DNA and reduce viscosity), then clarified (20,000 g, 15 min, 4°C). Protein concentration was determined using the Pierce™ 660 nm Protein Assay (Thermo Scientific), and equal protein was resolved by SDS–PAGE and transferred to PVDF. Membranes were probed for SREBP2 and a loading control (β-actin).

Band intensities were quantified by densitometry (ImageJ/Fiji), normalised to the loading control, and expressed as percentage of the 0 h value. Data are mean ± s.d. from n = 3 independent experiments.

**GFP pull-down**

Cells expressing endogenous or GFP-tagged APC were harvested at ~80% confluence, washed twice with ice-cold PBS, and resuspended in lysis buffer (25 mM Tris-HCl pH 7.5, 110 mM NaCl, 1.5 mM $MgCl_2$, 1mM TCEP) supplemented with EDTA-free protease inhibitors. Cell disruption was performed by nitrogen cavitation using a Parr cell disruption vessel (800 psi $N_2$, 20 min, 4°C), followed by rapid depressurization. Lysates were clarified by centrifugation (20,000 g, 15 min, 4°C), and the supernatant was collected for immunoprecipitation.

For affinity capture, clarified lysates were incubated overnight at 4 °C with pre-equilibrated GFP-Selector magnetic beads (NanoTag Biotechnologies, cat. no. N0315-

L) or control Selector Control magnetic beads (NanoTag Biotechnologies, cat. no. N0015-S), using 10–25 µL bead slurry per mg of lysate. Beads were gently rotated end-over-end during incubation.

Beads were separated magnetically and washed six times with wash buffer (25 mM Tris-HCl pH 7.5, 110 mM NaCl, 1.5 mM $MgCl_2$, 1 mM TCEP) at 4 °C. Bound proteins were eluted directly by adding 4× SDS sample buffer, vortexing briefly, and heating at 80°C for 5 min. Input, flow-through, final wash, and eluate fractions were collected for SDS–PAGE and immunoblotting.

**Generation and validation of APC-, AXIN1-, and β-catenin-knockout cell lines**

RKO and 293T cells were cultured in DMEM/F-12 medium (Gibco) supplemented with 10% FBS (Corning).

The HiFi Cas9 expression plasmid was generated by introducing a R691A mutation into the Cas9 coding sequence in pET-Cas9-NLS-6xHis (Addgene plasmid # 62933; Vakulskas et al., 2018). HiFi Cas9 protein was expressed in Rosetta™(DE3)pLysS Competent Cells (Novagen) and purified as described elsewhere (Zuris et al., 2015). The AsCas12a expression plasmid, generated by deleting the MBP sequence from plasmid pDEST-hisMBP-AsCpf1-EC (Addgene plasmid # 79007), was transformed into Rosetta(DE3)pLysS Competent Cells (Novagen) for expression. AsCas12a protein was purified as described elsewhere (Hur et al., 2016).

To generate APC knockout RKO cells, the sgRNA (target sequence GTCTTAGTGTAATACTGTAG) was generated using GeneArt Precision gRNA Synthesis Kit (Thermo Fisher Scientific). Transfection was performed with the Neon Transfection System (Thermo Fisher Scientific). Briefly, 0.6 µg sgRNA was incubated with 3 µg HiFi Cas9 protein for 10 minutes at room temperature and electroporated into 2x105 RKO cells.

To generate β-catenin knockout 293T cells, 80 pmol of Alt-R CRISPR-Cas12a crRNA (IDT) was incubated with 63 pmol of AsCas12a protein for 10 minutes at room temperature and electroporated into 2x105 cells along with 39 pmol of Alt-R Cpf1 Electroporation Enhancer (IDT).

The sgRNA target sequences for β-catenin-KO, GCTGAACCATCACAGATGCTGAAA.

Transfected cells were sorted as single cells into 96-cell plates. Knockouts were identified by Illumina MiSeq sequencing and further confirmed by Western blot.

**Extended Data Figure 1. Recombinant APC and AXIN1 purification and in vitro β-catenin degradation assays.**

**a,** Coomassie-stained SDS–PAGE gel showing sequential fractions from APC purification (input, flow-through, wash, and elution).

**b,** Western blot of purified APC using an APC-specific antibody confirming full-length protein.

**c,** Coomassie-stained SDS–PAGE gel showing AXIN1 purification (input, flow-through, first and second wash, and elution).

**d,** In vitro lysate-based β-catenin degradation assay using APC knockout (KO) extracts supplemented with increasing concentrations of purified APC (58.4, 116, and 175 nM). $^{35}$S-labelled β-catenin was used as a substrate.

**e,** AXIN1-dependent in vitro degradation of wild-type and phosphorylation-deficient (S33A, S37A, T41A, S45A) β-catenin in Axin1 KO extracts supplemented with increasing concentrations of purified AXIN1 (100, 200, 300, and 400 nM). $^{35}$S-labelled β-catenin was used as a substrate.

**f,** In vitro β-catenin degradation in HEK293T extracts supplemented with increasing concentrations (400 and 500 µM) of UbOK (lysine-less ubiquitin).

**Extended Data Figure 2. APC-dependent regulation of SREBP2 stability.**

**a,** Workflow of APC substrate identification from pooled TF screening.
A pooled in vitro degradation screen of 1,342 transcription factors identified 126 preliminary candidates upon repeat testing. From these**,** 44 potential candidate TFs**—**not confirmed positives—were individually tested in cell-extract degradation assays with 100 nM recombinant APC**.** Dose–response assays using increasing APC and AXIN1 concentrations have validated 4 substrates to date (SREBF1, SREBF2, MECP2, and CTNNB1).

**b,** In vitro degradation of $^{35}$S-labelled SREBP2 in Expi293 cell extracts with or without proteasome inhibitor. GAPDH served as a control for nonspecific turnover.

**c,** In vitro SREBP2 degradation in the presence of purified wild-type ubiquitin (Ub) or lysine-less ubiquitin (Ub0K) at increasing concentrations (100, 200, 300 µM). The inset shows the same gel at higher contrast to visualize the ubiquitin ladder.

**d,** Western blot analysis of SREBP2 in RKO and RKO APC knockout (KO) cells. Cells were pre-cultured in sterol-depleted medium and then switched to complete medium for 4 h to induce SREBP2 processing.

**e,** SREBP2 levels in HEK293T, HEK293T Axin1 KO, and HEK293T APC KO cells after 16 h cholesterol depletion followed by 4 h in complete medium.

**f,** HCT116 cells stably expressing doxycycline-inducible shRNA targeting APC show dose-dependent reduction of APC and accumulation of full-length SREBP2. Cells were treated with 0, 2, 4, or 6 µg/ml doxycycline.

**g,** Cycloheximide pulse-chase assay of SREBP2 turnover in RKO and RKO APC KO

cells. Cells were cholesterol-depleted for 16 h, then released into serum-containing medium with cycloheximide and harvested sequentially (0, 0.5, 1, 2, and 4 h). Representative blots (left) and quantification of full-length SREBP2 decay (right, mean ± s.d. from three independent experiments). SREBP2/actin ratios at each time point was normalized to the ratios at $t_0$.
**h,** Endogenous APC pull-down from APC–GFP knock-in cells. Lysates prepared by Parr bomb disruption were incubated with control or GFP magnetic beads, and bound proteins were immunoblotted for APC and SREBP2.

**Extended Data Figure 3. SREBP2 responsive protein expression in RKO versus RKO APC-KO cells.**
Cells were cholesterol-deprived for 16 h, then shifted to complete medium for 4 h. Whole-cell lysates (RIPA) were immunoblotted for LDLR, HMGCR, HMGCS, FDFT1, FDPS and MVK; β-actin is the loading control.

**Extended Data Figure 4. Activation of SREBP2 by Wnt requires APC and DVL.**
**a,** HEK293T and HEK293T APC knockout (KO) cells were cultured in cholesterol-depleted medium for 16 h, then switched to complete medium supplemented with recombinant Wnt Surrogate-Fc (50 ng/ml) and R-spondin (50 ng/ml). Cells were harvested at the indicated time points (0, 0.5, 1, 2, and 4 h), lysed in RIPA buffer, and immunoblotted for full-length and cleaved SREBP2; β-actin served as loading control.
**b,** HEK293T *DVL* KO cells treated in parallel under the same cholesterol-depletion and Wnt Surrogate-Fc/R-spondin stimulation conditions and analyzed for SREBP2 processing as in (a).
**c,** HEK293T *FZD* KO cells treated with cholesterol-depletion and Wnt Surrogate-Fc/R-spondin stimulation conditions as above and analyzed for SREBP2 processing as in (a&b).

**Extended Data Figure 5. FBXW7 regulates SREBP2 levels independently of GSK3β/CK1α activity.**
**a,** Validation of FBXW7 knockdown in HEK293T cells transfected with control or FBXW7 siRNA by immunoblotting.
**b,** Western blot analysis of the same cells shows increased full-length SREBP2 upon FBXW7 knockdown.
**c,** RKO cells treated with 2 μM LY2090314 (GSK3β inhibitor) or 2 μM SJ3149 (CK1α degrader) show the expected accumulation of β-catenin but no appreciable change in SREBP2 protein abundance. β-actin served as a loading control.
**d,** RKO cells treated with 1 or 2 μM LY2090314 (GSK3β inhibitor) show increased accumulation of SREBP1 protein abundance. β-actin was used as a loading control.

**Extended Data Figure 6. Purification of C-terminally truncated APC mutants.**
Western blot analysis of purified recombinant APC proteins harboring patient-relevant

C-terminal truncating mutations (T1556*, Q1338*, and S811*). Each lane shows affinity-purified APC variants following StrepII-based purification and detection with APC antibody, confirming production of the expected truncated proteins.
